# Sodium Aurothiomalate Induces Ferroptosis by Targeting GPX4 via Gold-Dependent Thiomalate Covalent Modification

**DOI:** 10.1101/2025.11.13.687868

**Authors:** Martín Hugo, Lissy Z. F. Gross, Andresa Messias, Lucia Alcober-Boquet, Thibaut Vignane, Biplab Ghosh, Nesrine Aroua, Lora Denson, Susana Delgado-Martín, Kamini Kaushal, Umut Yildiz, Sebastian Müller, Antonio Martínez-Ruiz, Darío A. Estrin, Hellmut G. Augustin, Raphaël Rodriguez, Ricardo M. Biondi, Andreas Trumpp, Ashok Kumar Jayavelu, Qing Cheng, Manuel Etzkorn, Carsten Berndt, Elias S.J. Arnér, Milos R. Filipovic, Santiago Di Lella, Daniel Pastor-Flores, Hamed Alborzinia

## Abstract

Ferroptosis, an iron-dependent form of oxidative cell death, is predominantly regulated by glutathione peroxidase 4 (GPX4), making it a promising target for cancer therapy. However, the majority of GPX4 inhibitors, most of which contain a chloroacetamide moiety such as RSL3, are limited by poor pharmacokinetic properties and off-target effects, hindering their preclinical translation. Utilizing a range of interdisciplinary methodologies, we show that sodium aurothiomalate (ATM), a drug approved by many agencies, induces ferroptosis by covalently targeting GPX4 via formation of a selenenylsulfide bond. In preclinical models of neuroblastoma and acute myeloid leukemia (AML), ATM combined with ferric ammonium citrate (FAC) yields a synergistic effect, resulting in a significant reduction in tumor growth. Mechanistically, ATM disrupts GPX4 activity by covalently binding thiomalate to the active site selenocysteine, while modification of specific cysteine residues leads to destabilization of the protein and impaired binding to phospholipids. We propose that these covalent modifications are achieved through a unique reaction mechanism, in which the gold component of ATM acts as a thiol-masking carrier and is only transiently present, being subsequently displaced and allowing the reaction of the thiomalate moiety with the target selenocysteine or cysteine. Our data lay the foundation for development of novel, drug-like thiol-based GPX4 inhibitors.

## Main

The regulation of ferroptosis, a form of cell death characterized by phospholipid peroxidation, is critically dependent on glutathione peroxidase 4 (GPX4). By utilizing glutathione (GSH) as a substrate and through the selenocysteine residue (Sec46, U46) from the catalytic site, GPX4 reduces and suppresses the propagation of lipid peroxidation, thereby preventing the onset of ferroptosis^1^. Recent studies revealed an increased dependence on GPX4 in certain cancer contexts, notably in drug-tolerant persister (DTP) cancer cells, as well as certain cancer types such as *MYCN*-amplified neuroblastoma. This dependency creates a selective vulnerability that can be exploited through its pharmacological inhibition for targeted therapies^2–8^.

However, the development of selective GPX4 inhibitors suitable for preclinical translation remains a significant obstacle, despite the promising therapeutic potential. Although existing inhibitors like RSL3 and ML210 show strong efficacy in cell-based assays, they suffer from poor pharmacokinetics, limiting their effectiveness in *in vivo* settings^9^. Furthermore, recent cell-free assays, enabled by advanced production methods for selenoproteins such as GPX4, which ensure robust selenocysteine incorporation^10,11^, have raised doubts about the specificity of these inhibitors. These studies demonstrated that while RSL3 and ML210 fail to inhibit the activity of purified GPX4, they target instead human selenoprotein thioredoxin reductase 1 (TXNRD1)^12^, as well as additional components of the selenoprotein synthesis machinery^13^. Moreover, structural studies have further expanded our understanding of GPX4 regulation, identifying an allosteric binding site at cysteine 66 (C66) and a phospholipid-binding cationic region around cysteine 148 (C148), reported as essential for the enzyme’s interaction with the phospholipid substrate^14,15^. In the drug discovery field, there is still great interest in covalent inhibitors^16^ and specifically for GPX4, most current efforts still predominantly rely on covalent modification of the active-site selenocysteine^17^.

In this study, we demonstrate that aurothiomalate (ATM), a gold-derived compound already approved as a drug, offers unique chemical properties for therapeutic exploration through ferroptosis induction by targeting GPX4. Indeed, *in vivo* experiments in neuroblastoma and acute myeloid leukemia (AML) xenograft models revealed that ATM, in combination with ferric ammonium citrate (FAC), synergistically suppresses tumor growth, supporting its potential application in GPX4-dependent cancers. Beyond its therapeutic implications, the effect of ATM on GPX4 offers important insights into the enzyme’s regulatory mechanisms. Our study reveals a unique mode of ATM-mediated inhibition, in which ATM covalently modifies GPX4 at three distinct sites: the active-site U46, as well as C66 and C148. These covalent modifications result in enzymatic inhibition, conformational destabilization, and disruption of phospholipid binding, collectively leading to the induction of ferroptosis. Furthermore, our data elucidate a novel mechanism in which gold serves solely as a thiol-protector carrier for the bioactive thiomalate moiety that is responsible of forming the covalent GPX4 modifications. However, the absence or presence of gold in the compound can determine the drug-like properties as well as the target selectivity of the thiomalate functional group.

## RESULTS

### The gold-containing compound aurothiomalate triggers lipid peroxidation and ferroptosis

Elevated levels of free iron can cause cellular damage by catalyzing Fenton-mediated hydroxyl radical (^•^OH) production, a key driver of oxidative damage^18^ linked to ferroptosis, a unique form of cell death characterized by its dependence on iron as a defining hallmark^19^. However, recent studies have suggested that transition metals, such as gold, may enhance sensitivity to ferroptosis^20,21^. To investigate this, we utilized *MYCN*-amplified SK-N-DZ neuroblastoma cells, which we had previously identified as highly dependent on the key ferroptosis-regulating enzyme GPX4^6^. We first treated these cells with ferric iron (III) ammonium citrate (FAC) or chloro(dimethylsulfide)gold(I), to assess their response to iron and gold, and to determine whether they undergo ferroptosis. As expected, iron significantly reduced cell viability, an effect that was fully reversed by ferrostatin-1 (Fer-1), a specific ferroptosis inhibitor (Fig. 1a). Interestingly, not only FAC but also the gold(I) compound triggered cell death, which could be partially rescued with Fer-1, confirming the role of ferroptosis as a mechanism of cell death (Fig. 1b). Furthermore, the tested gold (I)-containing compound induced lipid peroxidation, a key driver of ferroptosis, which was effectively prevented by co-treatment with Fer-1 (Fig. 1c). To further assess the ability of gold-containing salts to trigger ferroptosis, we treated SK-N-DZ cells with two well-known gold-containing compounds, auranofin and aurothiomalate (ATM). Auranofin has been proposed to trigger ferroptosis; however, this remains a topic of debate as multiple studies report conflicting evidence^22^. We observed that while Fer-1 fully rescued cells from ATM-induced cell death, it did not provide protection against auranofin (Fig. 1d,e). This finding highlights the fact that although thioredoxin reductase 1 (TXNRD1) is proposed as the primary target for both gold(I) compounds^23^, ATM can induce ferroptosis through additional regulatory targets beyond TXNRD1. In line with this observation, previously published TXNRD1 inhibitors, such as TRi-1^24^ and DKFZ-682^25^, lack ferroptosis-inducing capability, as no cell rescue was observed when SK-N-DZ cells were co-treated with the ferroptosis inhibitor Fer-1 (Supplementary Fig. 1a,b). To evaluate the overall oxidative stress-inducing effects of ATM and auranofin, we treated cells for 24 hours with a sublethal dose of the compounds and analyzed levels of glutathione (GSH), the most abundant low molecular thiol-containing molecule in cells, which is typically oxidized upon generalized oxidative stress. While ATM did not alter intracellular GSH levels or the GSH/GSSG ratio at the examined time point, auranofin significantly reduced the GSH/GSSG ratio (Supplementary Fig. 1c–e). Moreover, we observed that similar to the commonly used ferroptosis inducing compound RSL3 and unlike auranofin, ATM triggers lipid peroxidation (Fig. 1f), which is inhibited by Fer-1 and the iron chelator deferoxamine (DFO). Consistent with this, combination treatment with ATM and sublethal doses of RSL3 as well as with a ferroptosis suppressor protein 1 (FSP1, also known as AIFM2, an enzyme that protects cells from ferroptosis independently of GPX4^26,27^) inhibitor, resulted in a strong synergistic effect (Supplementary Fig. 1f,g). This suggests complementary mechanisms of action between ATM and RSL3, with ATM leveraging unique properties as a ferroptosis inducer that distinguish it from other gold compounds tested here.

**Figure 1:**
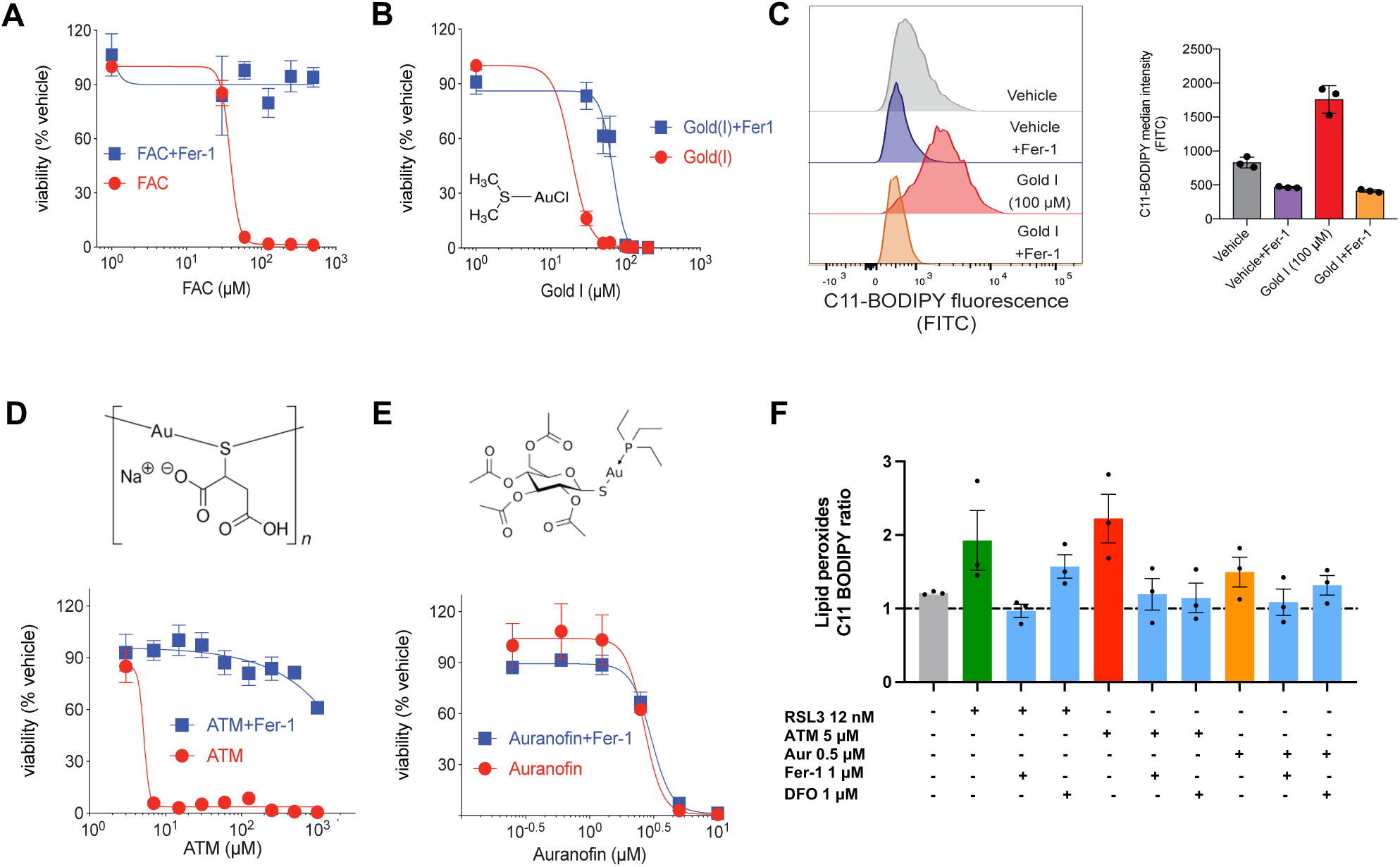
Induction of ferroptosis by gold and gold-containing compounds: a, b, SK-N-DZ cells treated with ferric ammonium citrate (FAC) and chloro-(dimethylsulfide)-gold(I) (Gold (I)) in the presence or absence of 5 μM ferroptosis inhibitor ferrostatin-1 (Fer1). Cell viability was measured after 3 days of treatment. c, Lipid peroxidation analysis upon treatment of SK-N-DZ cells with 100 µM of chloro-(dimethylsulfide)-gold(I) in the presence or absence of Fer1 using C11-BODIPY staining. Analysis was done after 1 day of treatment. d–e, SK-N-DZ cells treated with aurothiomalate (ATM) and auranofin (structures shown), for 3 days in the presence or absence of Fer1. f, Lipid peroxidation analysis upon treatment of SK-N-DZ cells with RSL3, ATM, and auranofin (Aur) in the presence or absence of 1 μM Fer1 and the iron-chelator deferoxamine (DFO) as ferroptosis inhibitors. Analysis was done after 24 h of treatment by using C11-BODIPY staining in 96-well plate; data are expressed as media ± sem of the ratio at 24 h with respect to the value at time.

### Selenium and modulation of ferroptosis regulators rescue ATM-induced ferroptosis

We further investigated the effects of ATM in the presence of various ferroptosis inhibitors, including the iron chelators ciclopirox olamine (CPX) and deferoxamine (DFO), as well as the lipoxygenase pathway inhibitor Zileuton, which has been shown to prevent ferroptosis^28^. Each of these inhibitors markedly improved cell viability in ATM-treated SK-N-DZ cells, strongly supporting ferroptosis as the primary mechanism of cell death (Fig. 2a). Additionally, ATM induced ferroptosis in a panel of neuroblastoma cell lines (Supplementary Fig. 2a). Notably cells with higher *MYCN* expression (LAN-1, IMR-32, SK-N-DZ) were more sensitive to ATM treatment, whereas the *MYCN*-low cell line SK-N-FI and human fibroblasts IMR-90 were more resistant. This is consistent with our previous findings that *MYCN*-amplified neuroblastoma cells exhibit increased sensitivity to ferroptosis induction, particularly through GPX4 inhibition^6^. However, this pattern of increased sensitivity in *MYCN*-high cells to ATM was not observed when the cells were treated with auranofin (Supplementary Fig. 2b). To further elucidate the ferroptosis-inducing mechanism of ATM, we pretreated the cells with 100 nM of sodium selenite (Na₂SeO₃) before exposing them to ATM (Fig. 2b) given that selenium is an essential trace element required for the activity of selenoproteins, including GPX4. Cells treated with Na₂SeO₃ were fully protected from ATM-induced cytotoxicity, whereas this protective effect was not observed with auranofin treatment (Fig. 2b,c). Consistent with this observation, overexpression of GPX4, as well as of FSP1 (also known as apoptosis-inducing factor 2, AIFM2), an enzyme that protects cells from ferroptosis independently of GPX4 ^26,27^, led to resistance to ATM but provided only moderate protection against auranofin (Fig. 2d-f). This is in line with previous reports of how overexpression of GPX4^29^ and FSP1^26,27^ correlates with the resistance of cancer cell lines to ferroptosis inducers like RSL3. Moreover, overexpression of the SELENOP transporter LRP8 in the SK-N-DZ cell line rendered the cells resistant to both ATM and RSL3 (Supplementary Fig. 2c). This suggests that ATM functions as a ferroptosis-inducing compound and highlights the differences in the mechanisms of action between ATM and auranofin. Furthermore, these findings underscore the unique ferroptosis-modulating properties of ATM, likely involving the key selenoenzyme GPX4. Having established that ATM-dependent cell death is primarily induced via ferroptosis and can be rescued by selenium supplementation and modulation of ferroptosis regulators, we proceeded to perform global transcriptional and proteomic profiling to uncover the broader molecular pathways affected by ATM treatment.

**Figure 2:**
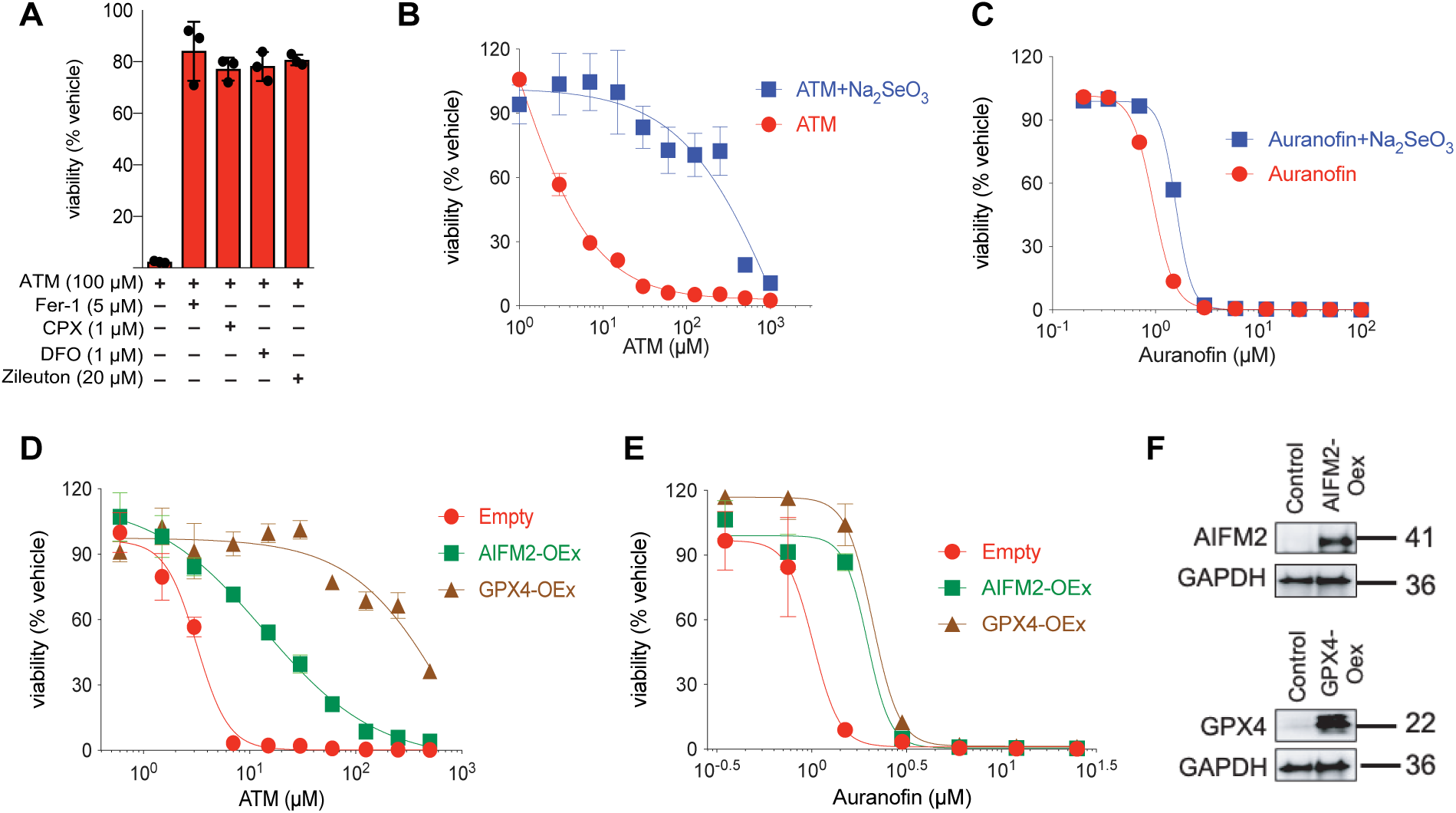
Modulation of ATM-induced ferroptosis by ferroptosis inhibitors. a, SK-N-DZ cells treated with 100 μM ATM in the presence or absence of ferroptosis inhibitors (5 μM Fer-1, 1 μM DFO, 1 μM CPX, or 20 μM Zileuton). b-c, Effect of sodium selenite supplementation on cell viability following treatment with ATM or auranofin. d-e, Survival of SK-N-DZ cells treated with ATM or auranofin with or without overexpression of GPX4 or AIFM2 (FSP1). f, Western blots showing successful overexpression of AIFM2 and GPX4. Data represents SEM; *p<0.05, **p<0.01.

### ATM modulates key ferroptosis regulatory pathways

To further understand how ATM regulates ferroptosis, we examined its cell-wide impact in SK-N-DZ neuroblastoma cells using transcriptomics and proteomics analyses. Treatment with ATM led to a significant activation of ferroptosis-associated pathways, as revealed by pathway analysis using the transcriptomics results (Fig. 3a,b). The top-regulated pathways included the NRF2 pathway, with significant upregulation of its key target gene NQO1 along with additional ferroptosis-related genes such as those involved in GSH biosynthesis (GCLC, GCLM), cysteine biosynthesis and uptake (CHAC1, SLC3A2, SLC7A11), metabolism, and transport of iron (FTH1, HMOX1, SLC40A1) in ATM-treated cells. In line with this, proteomics (Fig 3c-e and Supplementary Fig. 3a-c;e-j) and immunoblotting (Supplementary Fig. 3d) experiments validated marked alterations in the levels of ferroptosis-associated proteins at both 24 and 48 hours and with two different treatment concentrations. Both main oxidoreductases promoting cytosolic reducing pathways, GR and TXNRD1, were suppressed at 24 hours but displayed recovered levels at 48 hours (Supplementary Fig. 3g-h). ATM also induced changes at the protein level in key ferroptosis regulatory genes critical for selenium metabolism, such as an increase in LPR8, responsible for the cellular uptake of SELENOP^5^, as well as a pronounced reduction in GPX4 and in the recently identified selenium carrier protein PRDX6^30,31^ (Fig 3c-e). Moreover, peroxiredoxin 3 (PRDX3), recently reported as a ferroptosis marker^32^ was strongly upregulated at both time points tested (Fig 3 c-d). These findings further support the regulation of ferroptosis as a central mechanism underlying the cytotoxic effect of ATM. ATM treatment led to a concomitant increase in FSP1 (Supplementary Fig. 3d), indicative of a compensatory response aimed at limiting lipid oxidation via GSH-independent pathway, however insufficient, since the cells still undergo ferroptotic cell death. This highlights the disruption of the GPX4 system, which triggers a cascade of transcriptional and proteomic changes directly linked to oxidative damage and ferroptotic cell death, thereby highlighting therapeutic potential of ATM in ferroptosis-sensitive cancers.

**Figure 3:**
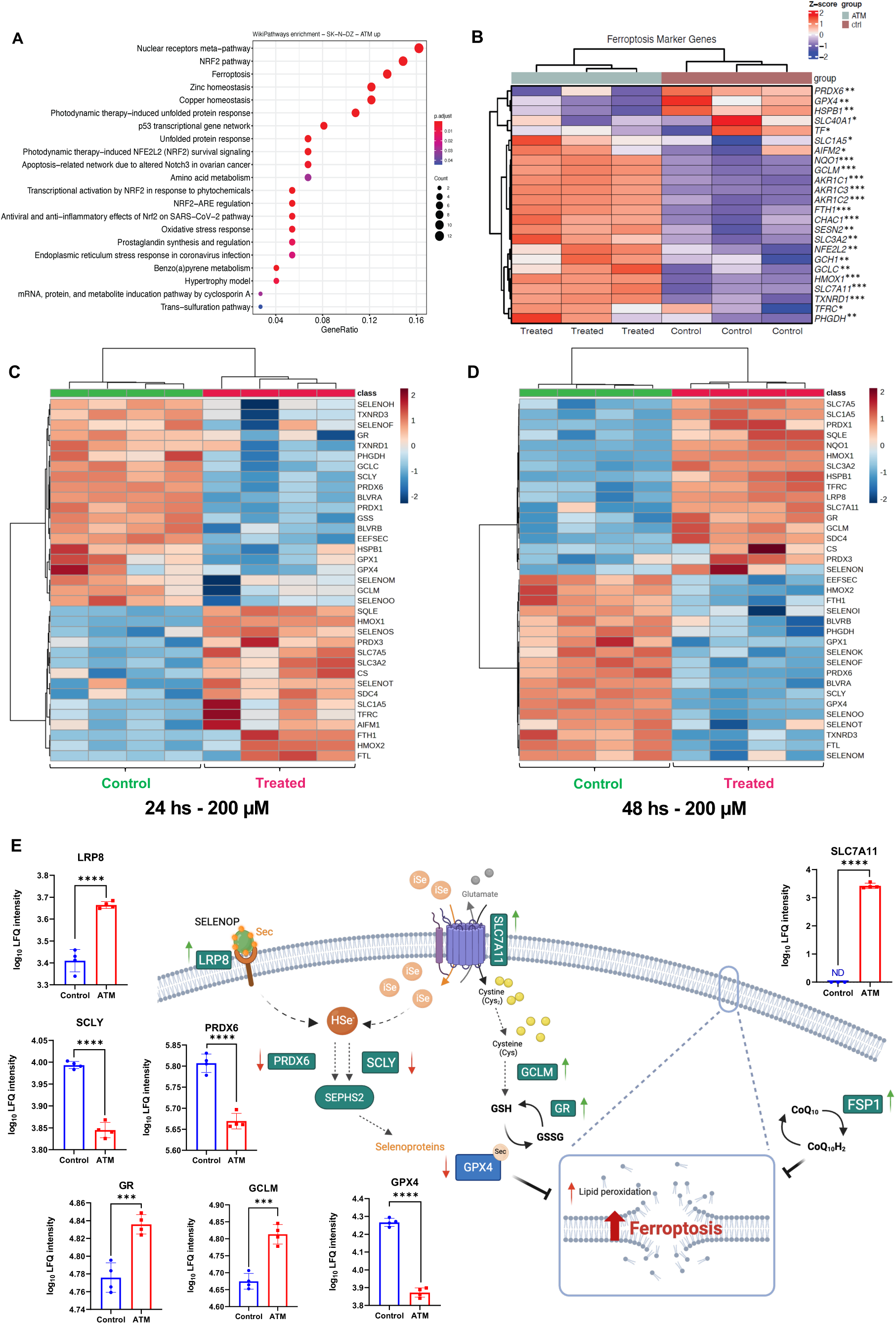
Gene and protein expression signature of ferroptosis upon ATM treatment: a, pathway analysis of SK-N-DZ cells treated with ATM for 24 hours. b, Transcriptional changes of ferroptosis markers of SK-N-DZ cells after 24 hours of treatment with ATM. ***P < 0.001; **P < 0.01. *P < 0.05. c-d, Protein level changes of ferroptosis markers in SK-N-DZ cells after 24 (c) or 48 hours (d) of treatment with media (left) or 200 μM ATM (right). e, Schematic representation of the proteins involved in selenium metabolism and ferroptotic pathways, whose expression is upregulated (green arrow) or downregulated (red arrow) upon 48 hours of treatment with ATM. Protein levels comparing control and ATM treatment after 48 hours are plotted as bars. ATM promotes degradation of GPX4, while other ferroptosis related proteins like LRP8, SLC7A11 and GR are increased. FSP1/AIFM2 was not detected in proteomics data, but immunoblotting (Supplementary Fig. 3d) also shows an upregulation. ****P < 0.0001 ***P < 0.001; **P < 0.01. *P < 0.05.

### Antitumor activity of ATM diminishes *in vivo* neuroblastoma and AML cell growth

Given that ATM induces ferroptosis in GPX4-dependent cellular systems, we next sought to evaluate its efficacy using *in vivo* models. However, its micromolar potency poses a challenge for its direct application in cancer therapeutics. To overcome this limitation, we explored a combination therapy approach to synergistically enhance ferroptosis-inducing potency of ATM. As a result, we observed that combining ATM with ferric ammonium citrate (FAC) significantly amplifies ferroptosis induction in our cell line model. The combined treatment of SK-N-DZ neuroblastoma cells with ATM and sublethal doses of FAC induced a pronounced synergistic cytotoxic effect, resulting in an approximately 20-fold reduction in the IC_50_ value of ATM. Notably, this synergy was specific for the ATM/FAC combination, as no similar effect was observed when FAC was combined with auranofin (Fig. 4a,b). FAC enhanced the potency of ATM at inducing ferroptosis, as evidenced by the protective effects of Fer-1 in cells co-treated with ATM and FAC (Fig. 4c). Based on this finding and given the potential of both compounds for preclinical applications, we performed *in vivo* experiments using two different cancer models: an orthotopic neuroblastoma model and an AML xenograft model (Fig. 4d). These experiments aimed to further evaluate the synergistic effect and assess whether this combination could be tolerated by the host. A 2-week treatment with ATM (80 mg/kg daily)^33^ or FAC (0.75 mg every other day) alone resulted in moderate reductions in tumor weight in the neuroblastoma model, with no significant impact on the overall health or weight of the mice (Supplementary Fig. 4a). However, combination therapy in this model yielded a significantly lower tumor weight, achieving approximately a 50% decrease, highlighting the enhanced efficacy of this synergistic approach (Fig. 4g,h). To evaluate whether ATM has reached the neuroblastoma tumor we performed ICP-MS analysis and observed a significant increase in intracellular gold content in tumor tissue isolated from mice treated with the ATM/FAC combination. In contrast, intracellular iron levels remained comparable to those of vehicle-treated mice (Fig. 4e,f). The observed iron-independent gold accumulation aligns with FAC’s role in promoting ferroptosis, by amplifying oxidative damage of cell membranes. We further evaluated ATM/FAC combination treatment in two distinct AML xenograft models, one using the unique cell line OCI AML 8227 previously generated by Dick *et al*.^34,35^ (Fig. 4i) and a second model based on a PDX derived from primary AML samples (Supplementary Fig. 4b,c). Combination therapy with ATM and FAC attenuated the number of AML blast cells in the bone marrow by approximately 50% using the PDX model, in agreement with the observations in the neuroblastoma model (Fig. 4g). The treatment of the AML PDX model with the ATM/FAC combination reduced tumor burden in both the spleen and bone marrow of recipient mice, though the reduction was statistically significant only in the spleen (Supplementary Fig. 4b,c). These findings highlight the therapeutic potential of combining ATM with FAC by exploiting the synergistic effects of GPX4 inhibition and FAC-induced oxidative damage and expand the application of ferroptosis-based therapies to the treatment of aggressive cancers, including neuroblastoma and AML.

**Figure 4:**
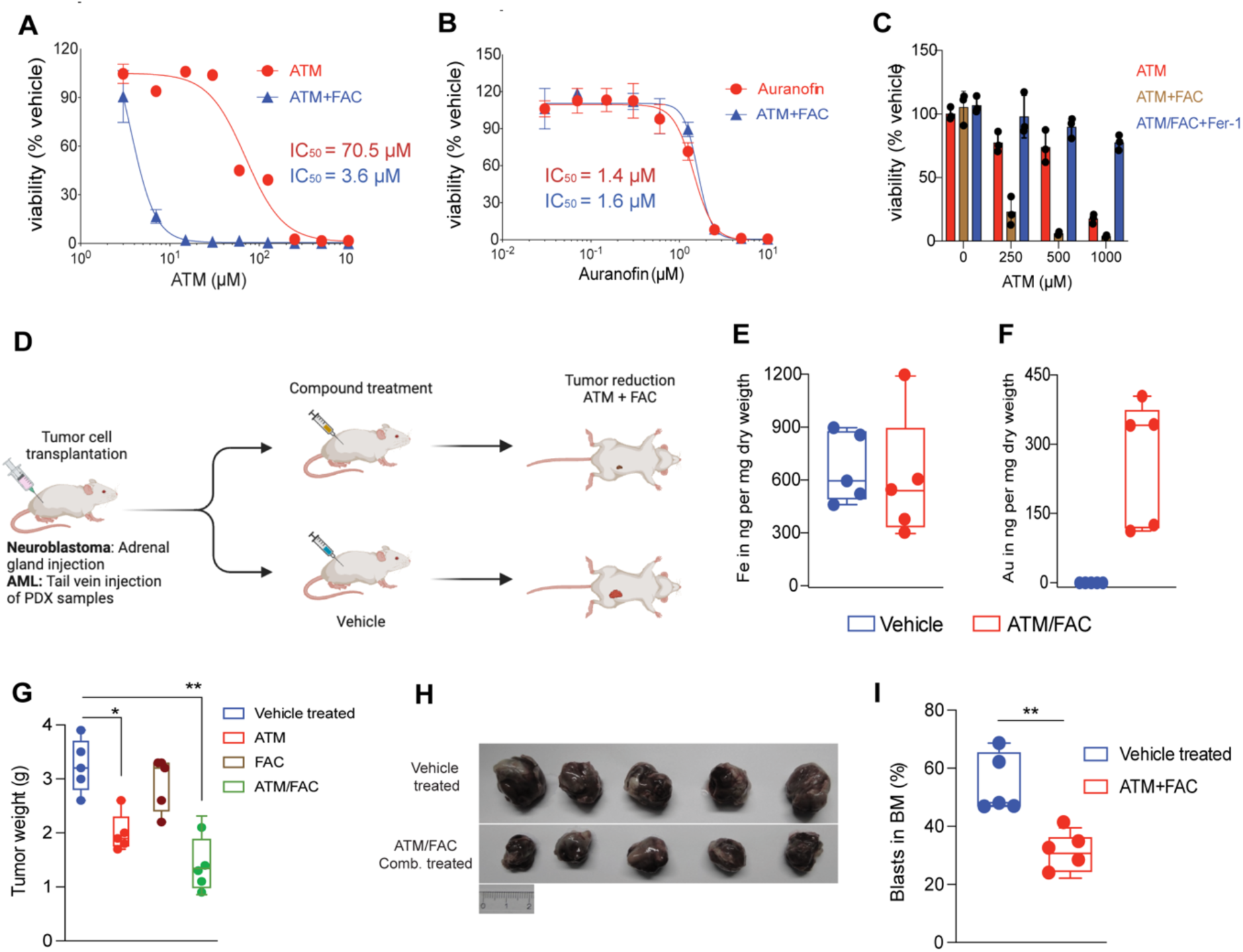
ATM/FAC combination attenuated orthotopic neuroblastoma tumor growth and the leukemic cells in AML model. a-b, SK-N-DZ cells treated with ATM or auranofin in the presence or absence of 40 µM of ferric ammonium citrate (FAC). c, SK-N-DZ cells co-treated with ATM and 40 µM of FAC in the presence or absence of Fer1. Cell viability was measured in all experiments after 3 days of treatment. d-i, *In vivo* experiments (n = 5) with ATM on two different cancer models: d, schematic representation of the *in vivo* experiments. e-f, Intracellular concentrations of gold and iron in neuroblastoma tumor cells isolated from mice treated with a combination of ATM and FAC were quantified using ICP-MS measurements. g-h, Orthotopic neuroblastoma was treated with individual ATM (80 mg/kg) or FAC, and a combination of both compounds for 2 weeks (5 days/week). g, Tumor weight of orthotopically implanted SK-N-DZ cells under the different treatment conditions (vehicle, blue; ATM, red; FAC, brown; ATM/FAC, green). h, Representative images of tumors in the control (blue) or ATM/FAC (green) treated groups. i, AML xenograft model was treated with ATM/FAC combination (80 mg/kg and 0.75 mg of FAC). Blasts percentage in bone marrow in both vehicle (blue) and ATM/FAC treated (red) groups.

### ATM Interacts with GPX4 and inhibits its activity

To elucidate the impact of ATM on GPX4 enzyme activity in greater detail, especially considering recent findings showing that GPX4 inhibitors effective in cellular systems do not necessarily inhibit the purified enzyme, we employed a GPX4/GSH reductase (GR) enzyme-coupled assay. This assay measures the activity of recombinant, selenocysteine (Sec)-containing GPX4 protein, following a previously described protocol^12^ (schematized in Fig. 5a).

**Figure 5:**
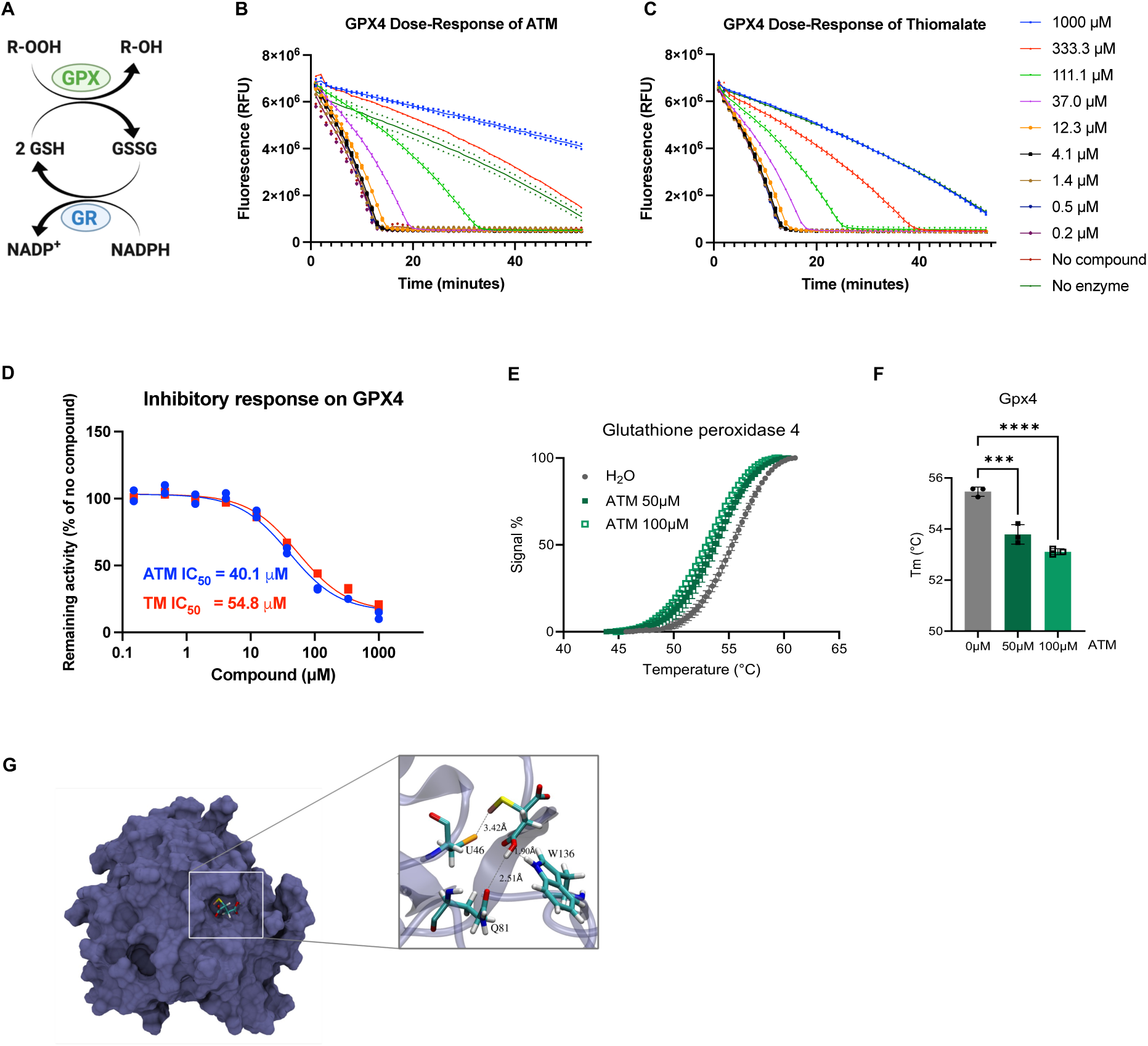
ATM targets the GPX/GR system: a, Schematic representation of the enzyme-coupled assay for the measurement of GPX activity. b-c, The activity of purified recombinant Sec-containing human GPX4 was measured using an enzyme coupled assay with glutathione reductase (GR), reduced glutathione (GSH) and NADPH as shown in (a), in the absence and presence of a wide range of ATM (b) or Thiomalate (c) concentrations (0.2-1000 µM). The reaction was initiated by the addition of 0.5 mM (final concentration in well) Cumene hydroperoxide to the reaction mixture and the NADPH fluorescence (λ_ex_ = 340 nm; λ_em_ = 450 nm) was monitored over time. d, Four-parameter dose-response curve fit to n = 2 technical replicates and IC_50_ calculations of ATM and Thiomalate. e-f, thermal shift assay with purified GPX4 at a final concentration of 1 μM and SYPRO orange dye, which binds to the hydrophobic patch of the protein once it is melted, resulting in a stronger fluorescence signal. g, Docking model for ATM in the region involving U46.

Our findings demonstrate that ATM inhibits GPX4 (Fig 5b) in a dose-dependent manner, impairing its enzymatic activity with an IC_50_ of 40.1 μM (Fig 5d). As a control, we observed that ATM caused a substantially less potent inhibition of GR activity as indicated by a higher IC_50_ of 187.8 μM (Supplementary Fig. 5c,h). Like GPX4, GPX1 (Supplementary Fig. 5a,g) activity was also inhibited by ATM. The observed higher potency is in accordance with the lower concentrations required for GPX1 degradation at a cellular level (Supplementary Fig. 3d). As expected for a gold-compound, ATM also inhibited TXNRD1 activity (Supplementary Fig. 5e,i). To further understand the mechanism of action of ATM, we also evaluated thiomalic acid, the corresponding gold-free analogue, across the different activity assays. Interestingly, thiomalic acid had a similar inhibitory potency to ATM with both GPX4 and GPX1 (Fig 5c-d, Supplementary Fig. 5 b,g), but no effect at all on GR (Supplementary Fig. 5d,h) and TXNRD1 (Supplementary Fig. 5f,i) activity. This correlates with our proteomics results, with the initial decrease of TXNRD1 and GR after 24 hours of treatment likely being due to ATM reactivity, while the protein levels are being restored or even increased after 48 hours. The latter could be due to compensatory mechanisms through NRF2 activation and the fact that the thiomalate moiety itself is not inhibiting these enzymes, assuming that thiomalate is formed over time from ATM (Supplementary Fig. 3g-h). Thus, GPX4 and GPX1 protein levels remain decreased after 48 hours, which would agree with the notion of thiomalate as well as ATM inhibiting these enzymes (Supplementary Fig 3e-f). Moreover, and in line with cell-based experiments and in agreement with earlier findings^12^, we did not observe any inhibition of GPX4 activity with auranofin, indicating once more the distinct modes of action of these two gold compounds (Supplementary Fig. 6b). We also did not observe any effect on GPX4 activity following incubation with RSL3, even at concentrations significantly higher than the IC_50_ derived from cell-based assays (Supplementary Fig. 6a), again in agreement with earlier results reported by Vučković *et al*. and Cheff *et al*.^12,36^. Together, these results highlight ATM’s distinct ability to target GPX4 and additional critical components of the cellular antioxidant defense system, thereby promoting ferroptosis.

To further validate GPX4 as a direct target of ATM, we conducted a thermal shift assay to assess changes in the thermostability of the enzyme upon interaction with ATM. Results revealed that ATM destabilizes GPX4, lowering its melting temperature and thus corroborating its direct binding and conformational perturbation of the enzyme (Fig. 5e,f). This destabilization could render the protein more susceptible to intracellular degradation, which is consistent with the dose-dependent decrease of GPX4 protein levels observed in our cellular assays (Fig. 3 c-e, Supplementary Fig. 3d).

Neither auranofin nor RSL3 affected the thermostability of GPX4 (Supplementary Fig. 6c,d), consistent with previous reports showing that RSL3 does not alter GPX4 melting temperature^12^. ATM also had no detectable effect on the thermal stability of GR (Supplementary Fig. 6e), suggesting a distinct mechanism of protein-ligand complex specific for GPX4. We next performed molecular docking simulations to gain further insight into the interaction between ATM and GPX4. The results suggest that ATM binds favorably within the active site of GPX4 (Fig. 5g). Residues Q81 and W136 were identified as critical for interacting with ATM’s carboxylic acid group, enabling the precise positioning of the gold-thiomalic moiety for covalent interaction with the active-site Sec residue (Fig. 5g).

Substitution of Sec with Cys significantly altered the predicted contacts and binding interactions in the model, underscoring ATM’s specificity for the active site U46 (Supplementary Fig. 7), which align with prior structural and mechanistic studies of selenocysteine-reactive ligands^37^. Overall, these findings suggest that ATM directly targets the activity and stability of GPX4, disrupting its enzymatic function and consequently inducing ferroptosis.

### Impact of ATM-dependent covalent modifications on structure and function of GPX4

To study the interaction between ATM and GPX4 more comprehensively, we examined whether ATM induces covalent modifications of the enzyme using wild-type GPX4 containing selenocysteine, as employed in activity and thermal shift assays. Mass spectrometry analyses revealed the formation of covalent bonds between the thiomalate moiety of ATM and GPX4 at three distinct residues: U46, forming a selenenylsulfide bond, and C66 and C148, forming disulfide bonds (Fig. 6a-c). Interestingly, we did not observe a covalent modification of either U46 or any of the cysteines by Au (to form RS(e)-Au, m/z +195.97) or the Au-thiomalate moiety (to from RS(e)-Au-CH (COOH)CH_2_COOH, m/z +345.10), indicating that the thiomalate moiety alone is responsible for the reactivity. Mechanistically, we propose ATM to form RS(e)-Au-SCH(COOH)CH_2_COOH as an intermediate, with gold subsequently catalyzing oxidation of thiols and selenium to form either a selenenylsulfide or a disulfide bond (Fig 6j,k). Quantitative peptide analysis revealed that U46 exhibited the highest reactivity, achieving maximal modification occurring at equimolar concentrations of ATM (Fig. 6d-f). This specificity is consistent with the higher reactivity of selenocysteine compared to cysteine, as supported by prior studies^38^. Molecular docking simulations further validated these findings, demonstrating the energetic favorability of ATM binding to GPX4 at all three sites. While only indicative for evaluating trend and not necessarily accurate in absolute values, the calculated binding energies for U46 (-3.1 kcal/mol), C66 (-2.3 kcal/mol), and C148 (-2.5 kcal/mol) confirmed U46 as the primary target (Fig. 6g-i). In order to assess the thermodynamical feasibility of the reaction of ATM with Cys and Sec residues, QM calculations were performed for model systems, and our results confirm that ATM preferentially reacts with selenium-containing residues presumably due to their higher acidity and lower bond dissociation energy than sulfur-containing residues (Supplementary Fig. 7 and Supplementary Table 1). These findings align with earlier reports describing the reactivity of selenium nucleophiles^38^. Beyond enzymatic activity, the modifications at C66 and C148 have additional regulatory implications. C66 resides within an allosteric modulation site of GPX4, while C148 is part of a cationic surface region critical for binding to negatively charged phospholipids^14,15^. Using Bio-Layer Interferometry, we confirmed that ATM impairs interaction of GPX4 with lipid nanodiscs containing negatively charged phospholipids, specifically 1-palmitoyl-2-oleoyl-sn-glycero-3-phosphoglycerol (POPG) (Fig. 6l). We also tested pure GPX4 incubation on lipid strips, where binding was observed mainly to phosphatidylglycerol and cardiolipin (also previously described by Cozza *et al.* ^15^), as well as low binding interactions with 3-sulfogalactosylceramide, phosphatidylinositol and sulfatide. Preincubation of GPX4 with ATM reduced those interactions (Supplementary Fig. 8). These findings highlight a unique functional aspect of ATM mode of action, wherein its thiomalate moiety modifies C148 and alters the lipid-binding properties of GPX4, making ATM an ideal compound for further chemical biology studies of GPX4 regulation. Molecular dynamics simulations of the GPX4-U46-TM adduct revealed a conformational destabilization, particularly in the 148-154 region (Fig. 6m-o), which is crucial for lipid binding^15^. Our structural perturbations data are consistent with our thermal stability assay results, which demonstrated an overall ATM-induced destabilization of GPX4 (Fig. 5e-f). These results imply that the inhibitory effects of ATM on GPX4 extend beyond the inhibition of enzymatic activity via binding of the thiomalate moiety to the U46 active site. Covalent modifications at C66 and C148 contribute to the structural destabilization and functional impairment of regulatory regions of GPX4. This multifaceted mechanism argues for ATM as a potential ferroptosis inducer through targeted disruption of GPX4 and presents a promising compound for studies focused on binding different regions of GPX4, which could offer valuable insight for researchers studying ferroptosis.

**Figure 6:**
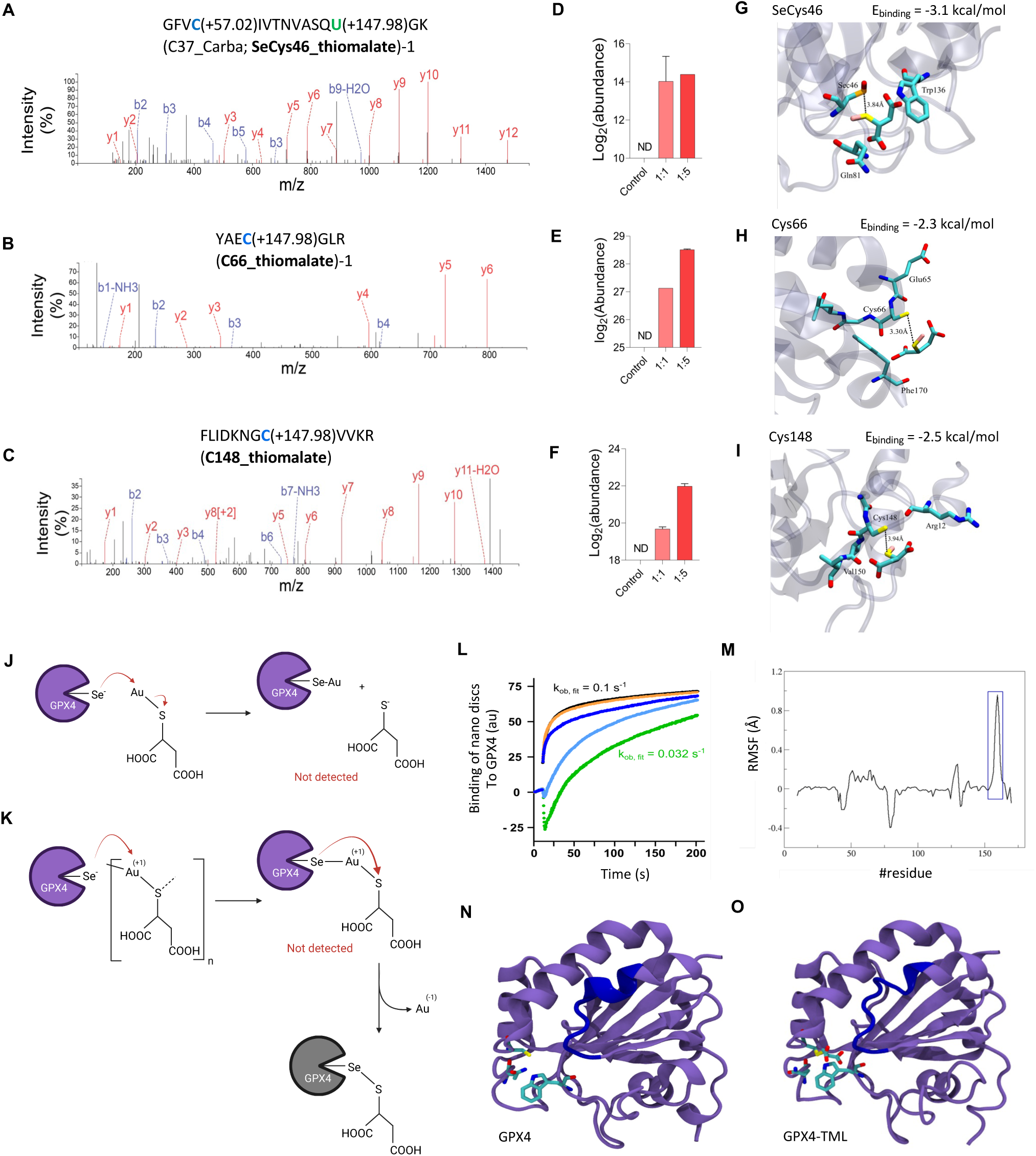
Covalent modification of GPX4 by ATM leads to loss of enzymatic function and destabilization. a-c, Peptide fragmentation spectra of identified covalent modification sites of free Sec (U, green) and Cys (C, blue) residues. d-f, Quantification of modified peptides shown in a-c, respectively, upon incubation of the protein with equimolar or 5x excess of ATM. g-i, Proposed best docking poses for ATM in three different sites: U46, C66 and C148. Binding energy values as derived from the docking analysis are shown as insets. j-k, Proposed reaction mechanisms between ATM and the (Se)Cys sites of GPX4. Only covalent bonds with the thiomalate moiety could be detected. l, Bio-Layer Interferometry following association of nanodiscs loaded with negatively charged 1-Palmitoyl-2-oleoyl-sn-glycero-3-phosphoglycerol (POPG) to GPX4, without and in the presence of 1 nM, 200 nM, 1 µM and 100 µM ATM (black, orange, blue, cyan and green lines, respectively). The observed rates of GPX4-POPG interaction (kobs) indicate no effects at the presence of ATM up to 1 nM and considerably slower binding at ATM concentrations of 1 µM and similarly at 100 µM.. m-o, Thiomalate covalently bound to Sec46 modifies GPX4 stability. m, Root mean square fluctuations (RMSF) difference between GPX4 in the free (n) and bound to TML (o) proposed states. While some regions experience minor differences in atomic fluctuations, a greater change is observed for the region between residues 152 and 158 (light blue), corresponding to a disordered region connecting the beta-strand comprising residues 148 to 154 and the alpha-helix comprising residues 159 and 171.

## DISCUSSION

This study establishes ATM, a compound approved by many pharmacological regulatory agencies, as a promising ferroptosis-inducing agent that targets GPX4, a critical regulator of cellular antioxidant and ferroptosis defenses. While ATM has been historically used to treat rheumatoid arthritis by modulating cyclooxygenase-2 and prostaglandin E2^39,40^, its potential as a ferroptosis inducer remained unclear until now. Numerous studies have demonstrated its efficacy in reducing joint inflammation and preventing disease progression, often with tolerable side effects. For instance, the molecular mechanisms of gold-based compounds like aurothiomalate highlighted their dual utility in anti-inflammatory and anticancer therapies^41^.

We here demonstrate that ATM targets GPX4 at the active site selenocysteine (U46), as well as other structural and functional regulatory regions. Notably, covalent modifications at C66 and C148 implicate ATM in altering the allosteric regulation, membrane-binding dynamics and protein stability of GPX4^14,15^. These interactions amplify inhibitory effects of ATM, further affecting GPX4 function and position ATM as a versatile ferroptosis inducer with a unique multi-site mechanism of GPX4 regulation, ultimately impairing GPX4 activity and stability. Interestingly, the observed destabilized region in our molecular dynamics simulation (Fig. 6m-o) also includes the R152 residue, mutations of which (e.g., R152H) are associated with Sedaghatian-type spondylometaphyseal dysplasia, a condition linked to GPX4 instability and degradation^42^.

Surprisingly, our mass spectrometry data indicates that only the thiomalate moiety remains covalently bound to its target residues. This observation is supported by the ability of thiomalic acid to inhibit GPX4 (but not GR or TXNRD1) activity with similar potency as ATM (Fig 5d; Supplementary Fig. 5 h-i), forming mixed selenenylsulfide and disulfide bonds that are not efficiently removed by the cellular enzymatic reducing systems. In contrast to ATM, when tested in cells, chloro-(dimethylsulfide)-gold(I) generated a minor induction of ferroptosis, while auranofin induced no ferroptosis at all (Fig. 1b,d,e). Therefore, ATM exhibits distinct pharmacological properties, where the gold component behaves as a carrier of the active thiomalate moiety through the formation of transient gold-containing intermediates with key regulatory selenocysteine and cysteine residues of its target proteins. This is in line with previous studies that have proposed mechanisms involving dynamic ligand exchange, where thiol-containing ligands, such as thiomalate or glutathione, displace gold from protein-gold adducts, potentially explaining the absence of gold in the observed covalent modifications^43,44^. Published crystallographic studies involving ATM are currently limited to only two structures of other protein targets (PDB 7YJR and 9H4V), both of which report the opposite effect, with binding of gold as the bioactive species to residues other than Cys or Sec, while thiomalate is absent due to previous release^45,46^. On the other hand, for PKCι it was assumed in the past that ATM forms a thiogold-cysteine adduct with the targeted Cys 69^47^. This underscores the novelty of the absence of gold in our proposed GPX4-targeting mechanism.

In addition to this, ATM also targets GPX1 (Supplementary Fig. 5g), which efficiently scavenges H₂O₂. Consequently, its inhibition by ATM leads to increased H₂O₂ levels, further amplifying oxidative damage^48^. Although ATM inhibits TXNRD1 activity as well (Supplementary Fig. 5 e,i), it displays distinct mechanistic features compared with the canonical TRXR1 inhibitor auranofin, not only because the latter fails to affect GPX4 activity (Supplementary Fig 6b-d), but also due to differences at a cellular level (Fig. 1d–e). The fact that previous exposure to sodium selenite protects SK-N-DZ cells against ATM but not auranofin (Fig. 2b–c) may be explained by the fact that GPX4 synthesis is more sensitive to perturbations in the selenocysteine metabolism than TXNRD1^49–51^. In line with this, our proteomics experiments show an initial decrease in TXNRD1 levels upon 24 hours of ATM treatment, that is later re-established to the control condition after 48 hours, likely due to NRF2 activation (Supplementary Fig 3. h). In contrast, GPX4 protein levels decrease consistently in both 24 and 48 hours (Supplementary Fig 3. e). A previous study that evaluates differences between auranofin and another gold-containing TXNRD1 inhibitor (aurothioglucose) demonstrates that ATM is not the first gold-compound with cytotoxicity and mechanistic differences to auranofin^52^. It should be noted that inhibition of TXNRD1 can trigger many downstream effects contributing to cell death^53^, and the TXNRD1-dependent TRP14 (also named TXNDC17) is the rate-limiting cytosolic reductase of cystine required for the use of cystine as a source of cysteine for GSH synthesis^54^. ATM therefore appears to inhibit several targets, all of which contribute to a general oxidative stress response that ultimately enhances inhibition of GPX4 and induction of ferroptosis. However, there seems to be a selectivity for GPX4 and GPX1 degradation at cellular levels (Supplementary Fig. 3e-f) in the long term, which aligns with the observed selective inhibition of these enzymes by thiomalate in the absence of gold (Fig. 5d; Supplementary Fig. 5g).

Combining ATM with ferric ammonium citrate presents a compelling strategy to overcome the moderate effect of ATM. The role of FAC in increasing intracellular labile iron pools amplifies oxidative stress via the Fenton reaction, intensifying lipid peroxidation and complementing GPX4 inhibition by ATM. This synergy highlights an innovative approach in ferroptosis-based therapy, leveraging a multi-site inhibition by ATM and the oxidative amplification induced by FAC. Moreover, this approach was effectively tested on drug-tolerant persister (DTP) cancer cells, which pose a significant challenge to durable cancer therapies and often rely on GPX4 to mitigate oxidative damage^2–6^. Combination therapy strategies address the limitations of single-agent treatments and enable the development of tailored regimens for specific cancer types. Notably, a Phase I clinical study demonstrated that such an approach is well-tolerated in cancer patients overexpressing protein kinase C iota (PKCι), including those with non-small-cell lung cancer, ovarian cancer, and pancreatic cancer^55^. The observed synergy also promotes the incorporation of other ferroptosis inducers targeting parallel antioxidant pathways, such as FSP1, to avoid compensating mechanisms and further enhance therapeutic efficacy^27^.

Moreover, thiol groups are rarely considered suitable warheads due to their chemical instability and metabolic liability^56^, with conventional GPX4 inhibitors usually possessing either a chloroacetamide (RSL3 and ML162) or a nitroisoxazole (ML210) group^17^. Our results prove that masking of the thiomalate active moiety by gold conjugation confers ATM its drug-like properties, which are critical for being safely and effectively used *in vivo*. Using different methods and experimental setups we demonstrated that ATM targets GPX4, providing a significant advantage over conventional GPX4 inhibitors, which were proven unstable and unsuccessful for preclinical experimentation. While these inhibitors are primarily assumed to interact with the active site, their efficacy remains controversial and is limited when tested on purified GPX4 or *in vitro* cell-free setups, with weak inhibitions in the micromolar range that are far from the well-studied low nanomolar concentrations sufficient for triggering ferroptosis in a cellular context^12,57,58^. Additionally, given that reduction of mixed disulfides requires the action of specific enzymes^59,60^, the covalent modification by thiomalate would most probably remain irreversible until the protein is degraded and further replaced by newly synthesized ones.

In conclusion, this study presents ATM as a direct GPX4 inhibitor and ferroptosis-inducing agent with a robust effect across different interdisciplinary experimental approaches, from mechanistic studies on pure protein to *in vivo* evaluation. Moreover, we analyzed its unique reaction mechanism, in which the gold is only present in transient intermediates and subsequently displaced for the final functional thiomalate-modification of the targeted Sec/Cys residues. Taken together, our findings lay the groundwork for the development of novel gold-conjugated thiol compounds with improved drug-like properties, capable of targeting not only GPX4 but also other selenocysteine-and cysteine-containing enzymes of interest.

## Acknowledgments

Martín Hugo is a Serra Húnter Fellow. Lissy Z. F. Gross is a DKFZ Postdoctoral Fellow. Work in the Alborzinia lab is supported by the support of DFG Priority Program SPP 2306 (AL 1533/5-1) and the Deutsche Jose Carreras Leukämie Stiftung (DJCLS 04 R/2024). SDM is a FPU fellow from the Spanish Government MICIU (fellowship FPU20/00995). AMR work was supported by grant PID2021-124688OB-I00 from the Spanish Government MICIU/AEI/10.13039/501100011033 cofunded by European Union ERDF (“A way of making Europe”). SM and RR acknowledge the Charles Defforey–Institut de France, European Research Council under the European Union’s Horizon 2020 research and innovation programme (grant agreement No 884754), Ligue Contre le Cancer (Equipe Labellisée), Institut National du Cancer (2022-1-PLBIO-06-ICR-1), Agence Nationale de la Recherche (ANR-23-CE44-0030). The research of QC and ESJA is supported by Karolinska Institutet, The Swedish Cancer Society (21 1463 Pj), The Swedish Research Council (2021-02214), The Cayman Biomedical Research Institute (CABRI), and The Hungarian National Research, Development and Innovation Office (projects 2022-2.1.1-NL-2022-00010, TKP2021-EGA-44 and K 146277). We thank Marcus Conrad for providing the Flag-strep-HA-tagged GPX4 expressing MEFs. CB acknowledges the DFG (417677437/GRK2576). HGA was supported by grants from the Deutsche Forschungsgemeinschaft (DFG) (project C05 within the CRC1366 “Vascular control of organ function” [project number 394046768] and the European Research Council ERC AdG “Angiomature” [project number: 787181”].

## Author contributions

M.H, D.P.F and H.A conceived the study. H.A. performed the majority of cell-based assays. Activity assays were performed by M.H. and L.Z.F.G with pure Sec-containing GPX4 provided by Q.C and E.A. Lipid peroxidation assay by spectrometric fluorescence was performed by S.D.M. under the supervision of A.M.R. Thermal stability assays were performed by L.A.B. under the supervision of R.M.B. Mass spectrometry was performed by T.B. under the supervision of M.R.F. Docking and molecular dynamics experiments were performed by A.M. under the supervision of D.A.E and S.D.L. In vivo experiments were performed by H.A. and N.A. with the support of A.T. Western blot was performed by K.K. Microarray assay was performed by U.Y. Proteomics experiments were performed by B.G. with the support of L.Z.F.G and under the supervision of H.G.A. and A.K.J. Bio-layer interferometry was performed by L.D under the supervision of M.E. and C.B. Lipid overlay assay was performed by L.Z.F.G. The manuscript was written by M.H., L.Z.F.G, D.P.F and H.A with support of S.D.L and input from all authors. All authors discussed the results and approved the final version of the manuscript.

## MATERIALS AND METHODS

### Chemicals

All chemicals used in this study were obtained from the following sources: ferrostatin-1, RSL3, and (1S,3R)-Methyl-2-(2-chloroacetyl)-2,3,4,9-tetrahydro-1 [4 (methoxycarbonyl)phenyl]-1H-pyrido[3,4-b]indole-3-carboxylate were from Tocris Bioscience; auranofin, Deferoxamine, tert-Butyl hydroperoxide, Gold(III) chloride, reduced glutathione (GSH), oxidized glutathione (GSSG), glutathione reductase (GR), and reduced nicotinamide adenine dinucleotide phosphate (NADPH) were from Sigma-Aldrich; Bodipy 581/591 C-11 was from Invitrogen; antibiotics, HEPES, DMEM, and DMEM Fluorobrite were from Gibco. The anti-GPX4 and anti-GPX1 antibodies were obtained from Abcam (1:1,000 dilution, catalog no. ab41787 and no. ab22604; Abcam), anti-LRP8 from LS Bio (1:1,000 dilution, catalog no. LS-C313670; LS Bio), anti-SLC7A11 from CST (1:1,000 dilution, xCT/SLC7A11 (D2M7A) Rabbit mAb, catalog no. 12691S; CST), AIFM2 from Proteintech (1:1,000 dilution, AIFM2/FSP1 Antibody, catalog no. 20886-1-AP; Proteintech), and beta-actin from Santa Cruz (1:5,000 dilution, beta actin antibody (C4), catalog no. sc-47778; Santa Cruz).

### Cell culture and analysis of cell viability

The human neuroblastoma cell line SK-N-DZ, obtained from ATCC, was cultured in DMEM medium supplemented with 10% FCS, 1× HEPES, and 100 U/ml penicillin/streptomycin at 37°C with 5% CO_2_. Identity verification was performed using SNP genotyping, and mycoplasma contamination testing was conducted by Eurofins. The effect of different compounds on cell viability was evaluated using the CellTiter-Glo (CTG) assay from Promega. To assess alterations in cell viability, 3,000 cells were plated in complete medium in 96-well plates (Greiner Bio One) 24 hours prior to treatment. Subsequently, cells were exposed to the specified concentrations of compounds for 72 hours. The CTG assay was employed to analyze cell viability according to the manufacturer’s guidelines.

### Generation of cells overexpressing genes

SK-N-DZ cells with constitutive expression of the CRISPR activation machinery were created by transducing wild-type cells with lentiviral particles containing dCas9-VP64 (the lenti dCas9VP64_Blast plasmid, Addgene #61425) at a multiplicity of infection (MOI) of approximately 0.5. Following recovery, cells were selected using Blasticidin treatment (20 μg/ml). To induce overexpression of GPX4 and AIFM2, individual gRNAs were cloned into the pXPR_502 vector (Addgene #96923) through restriction digestion of the respective lentivector with BsmBI (NEB, Cat. No. R0739). Oligonucleotides (from Sigma-Aldrich) containing the gRNA sequences and complementary overhangs were phosphorylated, annealed, and then inserted into the respective lentiviral delivery vector.

### Microarray Analysis

RNA extracted from SK-N-DZ cells treated for 24 hours (five vehicle-treated versus ATM treated) underwent analysis using the Affymetrix GeneChip Clariom S Assay. The microarray data (GSE192976) were processed using a modified version of a previously described pipeline^61^. In brief, raw CEL files were normalized using robust multichip average normalization, and two-group comparisons were conducted using the limma package v.3.46 (Bioconductor), employing an empirical Bayes test for differential expression. A false discovery rate (FDR) < 0.1 was considered statistically significant. Overrepresentation analysis was performed with clusterProfiler (v4.0.2) using terms retrieved from the Wikipathways database.

### In vivo mouse experiments

All experiments involving mice and experimental procedures were conducted in accordance with the guidelines set forth by the German Cancer Center Institute and were approved by the governmental review board of the state of Baden-Wuerttemberg, Regierungspraesidium Karlsruhe, under the authorization number G-176/19 and G205-22, following German legal regulations. The mouse strain utilized in the study was NOD.Cg-PrkdcscidIl2rgtm1Wjl/SzJ (NSG, JAX stock #005557). Female mice aged 3–4 months were employed for the experiments and were housed in individually ventilated cages under controlled temperature and humidity conditions. The cages were equipped with enrichment materials, including bedding. As previously outlined, orthotopic mouse models for neuroblastoma were established by transplanting 2 × 10^5^ SK-N-DZ cells into the right adrenal gland following surgical site preparation. The cells were suspended in a 1:1 (vol/vol) mixture of growth factor-reduced matrigel (Corning) and PBS. Subsequently, a total of 20 μl of this cell suspension was injected into mice under general anesthesia. Post-tumor cell transplantation, mice were closely monitored for evidence of tumor development using bioluminescent signal detection with an IVIS Spectrum Xenogen system. After the completion of tumor cell transplantation, treatment with ATM (80 mg/kg/day), either alone or in combination with iron citrate (1 mg -administered every second day), was initiated for two weeks. We observed a clear signal from the tumors 1 week after injecting 2 × 10^5^ SK-N-DZ cells. For AML xenograft model, cells were transplanted into sublethally irradiated (175 centi-Gray) NSG mice via intrafemoral injection. Upon detection of human engraftment, monitored by bone marrow aspiration, the mice received daily intraperitoneal injections of ATM and FAC, combination for a total of 10 days. At the experimental endpoint, all mice were sacrificed in accordance with European ethical protocols, and the spleen and bone marrow (from the tibia, femur, pelvis, and spine) were isolated and analyzed by flow cytometry. Throughout all animal experiments, the health of the animals was monitored daily. Mice were euthanized promptly upon meeting the criteria for humane endpoint, as defined in the experimental protocol. The sample size was calculated with the help of a biostatistician using R version 3.4.0. Assumptions for power analysis were as follows: *α* error, 5%; *β* error, 20%. Mice were randomized into treatment groups before treatment. If animals had to be sacrificed before the predefined endpoint (due to weight loss or other termination criteria), they were excluded from any downstream analyses. All animal experiments were blinded during experiments and follow-up assessments.

### Determination of intracellular lipid peroxidation levels

SK-N-DZ cells were seeded in black, opaque 96-well assay plates at a density of 5000 cells per well. Twenty-four hours later, the fluorescent probe Bodipy™ 581/591 C-11 (C11-BODIPY) was added to the wells at a concentration of 2 µM. Cells were incubated with the probe for 30 minutes, and then the medium was removed and replaced with DMEM Fluorobrite supplemented with 0,5% heat-inactivated FBS. This medium contained the respective treatment: RSL3 12 nM, aurothiomalate 5 µM or auranofin 0.5 µM. Each compound was added to the cells alone, or in combination with Ferrostatin-1 1 µM or deferoxamine 1µM. Cells were maintained for 24 hours in a standard cell incubator (95% air, 5% CO_2_ in gas phase, at 37 °C) protected from light. Wavelength spectra measurements were taken right after the addition of the treatment, as well as 24 hours later, using the microplate reader Clariostar Plus (BMG Labtech). Data are calculated as a ratio between the maximum fluorescence in the 520-530 nm range and the maximum fluorescence in the 590-600 nm range, blank corrected. Then, data are normalized to the initial measurement for each condition initial measurement. We also assessed lipid peroxidation using the BD FACSAria III cell sorter. For this, 10^5^ cells were plated in six-well plates. Prior to lipid peroxidation assessment, the medium was aspirated, and C11-BODIPY (with a wavelength of 581/591 nm) diluted in Hank’s Balanced Salt Solution (HBSS; Gibco) was added to the wells, reaching a final concentration of 4 μM. Following a 15-minute incubation period at 37°C in the tissue culture incubator, cells were gently harvested, and lipid peroxidation levels were promptly evaluated using a BD FACS Aria™ III cell sorter.

### Proteomics

**Proteomic sample preparation:** For protein extraction, the cell pellet was lysed with 100 μL of 1% SDC buffer, followed by boiling at 95 °C for 15 min and bioruptor for 20min with 30s pulses on and off for 20cycles. For tryptic digestion, Trypsin/LysC (1:100) was added and incubated for 16 hours at 37 °C at 1400rpm. The digestion was stopped by adding 5X volume of Isopropanol containing 1% TFA. Three layers of Styroldivinylbenzol-Reversed Phase Sulfonate (SDB-RPS) stagetips were equilibrated with 30%MeOH/1%TFA and 0.2% TFA. 30ug was loaded onto the stagetips and washed with isopropanol/1% TFA and 0.2% TFA. Peptides were eluted with 60μL of 80% acetonitrile (ACN)/1.25% ammonium hydroxide. The peptides were dried at 45°C for 90 min using a speed vacuum concentrator and then resuspended in buffer A* (2% ACN, 0.1% TFA). The concentration of samples was quantified using a Nanodrop, and the samples were stored at -20°C. **Data-independent acquisition (DIA) measurement:** Peptides (200 ng) were injected onto a nanoElute system coupled to a TIMS TOF Pro mass spectrometer (Bruker Daltonics) via a CaptiveSpray nano-electrospray ion source. Separation was performed on an IonOpticks Aurora 25 cm column using a binary solvent system comprising buffer A (0.1% formic acid, 2% acetonitrile) and buffer B (0.1% formic acid, 99.9% acetonitrile). For peptide separation, a 120-minute gradient was applied, starting at 2% buffer B and increasing to 12% over 60 minutes, 20% over 30 minutes, 30% over 10 minutes, and 85% over 10 minutes, followed by a 10-minute hold at 85% buffer B, with a flow rate of 0.3 µL/min. TIMS calibration employed Agilent ESI-Low Tuning Mix ions (m/z 622, 922, 1222) with known reduced ion mobility coefficients (1/K0). Data were acquired in dia-PASEF mode across m/z 100–1700 and 1/K0 0.65–1.40. The collision energy was set by linear interpolation between 80 eV at an inverse reduced mobility (1/K0) of 1.60 versus/cm2 and 20 eV at 0.6 versus/cm2. TIMS parameters were: scan range 100–1700 m/z, ramp time 100 ms, 100% duty cycle, cycle time 100 ms, and spectral acquisition rate of 9.43 Hz. **MS data processing:** The raw mass spectrometry data were analyzed using DIA-NN (v1.8.2)^62^ search engine with default parameters. Data were searched against the Homo sapiens UniProt reference proteome (entries from 7th March, 2021). ‘‘trypsin/ P’’ was specified as the protease, allowing up to one missed cleavage. ‘‘N-term M excision’’ and ‘‘C carbamidomethylation’’ were set as fixed modifications. A theoretical spectral library was constructed using deep learning algorithms, and a decoy database was included to calculate the false discovery rate (FDR) caused by random matches. The FDR for precursor identification was set at 1%. Precursor and protein FDR were both set to 1%; proteins must be identified with at least one peptide. **Bioinformatic analysis:** After DIA-NN analysis, the data set comprised the identified proteins, along with LFQ intensities. The data were filtered to remove features matched with contaminants. Following missing value imputation, the log transformed data was used for statistical analysis using MetaboAnalyst^63^ software. Heatmap representing the expression of ferroptosis regulators also obtained from statistical analysis in MetaboAnalyst. Metascape tool was used to obtain statistically enriched terms (GO/Kyoto Encyclopedia of Genes and Genomes pathway [KEGG] terms, Reactome and canonical pathways) for all differentially expressed proteins. All analyses were performed in four independent biological replicates.

### Activity assays

Recombinant Sec-containing enzymes were cloned, produced and purified as described previously^54^. GPX4 and GPX1 activity was assayed fluorometrically employing a coupled enzyme test in a 96-well plate following a similar protocol as the one described by Cheff *et al*^12^. Each well was loaded with 10 µL of 10x compound (ATM, thiomalic acid, RSL3, auranofin) to reach the indicated final concentrations. 90 µL of enzyme in Assay Buffer (0.1M Tris-HCl buffer, pH 7.4, 5 mM EDTA, 0.1% Triton X-100 and 0,01 % fatty acid-free BSA) were added in order to incubate the compounds with 200 nM of GPX4 or 80 nM of GPX1 for 30 min at room temperature. Then, 20 μL of master mix (final concentration in well 100 nM GR, 1 mM GSH, 0.5 mM NADPH in Assay buffer) were added. To start the reaction, 5 μL of Cumene hydroperoxide diluted in 50% EtOH was added for a final concentration of 0.5 mM. The decrease in NADPH fluorescence (λ_ex_ = 340 nm; λ_em_ = 450 nm) was monitored over time. For the GR counter assay, 2 nM of GR were incubated with the compounds of interest for 30 minutes at room temperature. To achieve this, 10 μL of 10x compound were added to 90 μL of enzyme in Assay buffer. After incubation, the reaction was started by dispensing 25 μL of GSSG and NADPH master mix in Assay Buffer for final concentrations of 1 mM GSSG and 0.4 mM NADPH. The decrease in NADPH fluorescence was recorded as for the GPX activity. For the TXNRD1 assay, each well was loaded with 10 µL of 10x compound (ATM or thiomalic acid) to reach the indicated final concentrations. 90 µL of enzyme in Assay Buffer (0.1M Tris-HCl buffer, pH 7.4, 5 mM EDTA, 0.1% Triton X-100 and 0,01 % fatty acid-free BSA) were added to incubate the compounds with 20 nM of TXNRD1 for 30 min at room temperature. After adding 5 µL NADPH, the reaction was triggered with 20 µL DTNB, with final concentrations in well of 0.3 mM NADPH and 2.5 mM DTNB. Thiomalic acid was freshly dissolved in water immediately prior to use in each experiment. Activity assays were performed in at least two independent experiments with consistent results. The figures shown represent one representative experiment including technical replicates.

### Temperature stability assay

Protein unfolding was monitored by the increase in the fluorescence of the fluorophor SYPRO Orange (Invitrogen) using a real-time PCR device (StepOnePlus, Applied Biosystems) following the protocol described^64^. Proteins were diluted in 10 mM HEPES (pH 7.5) buffer containing 200 mM NaCl. The reactions were performed in a final volume of 10 μL in 96-well PCR microtiter plates (Applied Biosystems) and contained 1 μM protein, 10 mM HEPES (pH 7.5), 200 mM NaCl, 1/1000 SYPRO Orange, and 2 mM reduced glutathione. ATM or water were added to this reaction mixture. The temperature gradient was performed in steps of 0.3°C in the range of 25°C to 85°C. To calculate the T*m* values, the data were exported to GraphPad Prism and the curves fitted to a Boltzmann sigmoidal equation with all R 2 > 0.998.

### LC/MS analysis of purified GPX4

1 µM of the purified protein was incubated either with 1 µM, or 5 µM of aurothiomalate for 1 hour at 37 °C in 50 mM ammonium bicarbonate. Reaction was then blocked by incubating the protein with 50 µM iodoacetamide. The protein was then subjected to tryptic digestion (Trypsin: Protein, 1: 20, w/w) for 20 h at 37 °C, 750 rpm. Peptides were desalted using Supelco desalting columns and vacuum dried. Samples were resuspended in 0.1 % trifluoroacetic acid (TFA) and peptide quality control was performed using an Ultimate 3000 Nano Ultra High-Pressure Chromatography (UPLC) system with a PepSwift Monolithic® Trap 200 µm * 5 mm (Thermo Fisher Scientific).

Peptides were analyzed by high-resolution LC-MS/MS using an Ultimate 3000 Nano Ultra High-Pressure Chromatography (UPLC) system (Thermo Fisher Scientific) coupled with an Q Exactive HF Mass Spectrometer via an EASY-spray (Thermo Fisher Scientific). Peptide separation was carried out with an Acclaim™ PepMap™ 100 C18 column (Thermo Fisher Scientific) using a 55 min linear gradient from 3 to 35 % of B (84 % Acetonitrile, 0.1 % Formic Acid) at a flow rate of 250 nL/min. The Orbitrap Eclipse™ was operated in a DDA mode, and MS1 survey scans were acquired from m/z 300 to 1,500 at a resolution of 60,000 using the Orbitrap mode. MS2 scans were carried out on top 15 ions with high-energy collision-induced dissociation (HCD) at 32 % using at a resolution of 15, 000.

Data were evaluated with PEAKS ONLINE software using following parameters: 15 ppm for precursor mass tolerance, 0.5 Da for fragment mass tolerance, specific tryptic digest, and a maximum of 3 missed cleavages. Following modifications were used as variable modification for peptide analysis: N-term acetylation (+42.010565 Da), methionine oxidation (+15.994915 Da). Carbamidomethylation (+57.021464 Da), gold (+195.958775 Da), aurothiomalate (+344.9496 Da), and thiomalate (+147.983 Da) on C were added as variable modification. For Selenocysteine, the fasta file with selenocysteine 46 replaced by cysteine was used and the mass difference between Se and S was included into the used modifications: Se-Carbamidomethylation (+104.965964 Da), Se-gold (+243.903275 Da), Se-aurothiomalate (+392.8941 Da), and Se-thiomalate (+195.9275 Da) on C were added as variable modification.

### Bio-Layer Interferometry

Bio-Layer Interferometry (BLI) was carried out by using the BLItzTM device from avantorTM and the high precision streptavidin Octet^®^ SAX biosensors. All steps were performed at room temperature using nanodiscs compromised of negatively charged 1-Palmitoyl-2-oleoyl-sn-glycero-3-phosphoglycerol (POPG) lipids to follow the association and dissociation with GPX4 monitored in real time. Prior to use, the biosensor was hydrated for 20 minutes in sodium phosphate buffer (20 mM sodium phosphate, 50 mM sodium chloride pH 7.4). After hydration, the initial baseline was measured for 30 seconds. Subsequently, 4 µl of protein lysate of MEFs expressing flag-strep-HA-tagged GPX4^49^ were loaded onto the drop holder of the BLI device and GPX4 was immobilized on the tip of the biosensor. Loading of the protein to the sensor was then measured for 300 seconds and the sensor placed back into sodium phosphate buffer for a second baseline measurement. Next, 4 µl of POPG nanodiscs without or with ATM (1 nM, 200 nM, 1 µM, 100 µM) were loaded onto the drop holder and association monitored for 180 seconds. Different dilutions of GPX4 were measured and a negative control was carried out by using buffer. Data were processed using baseline compensation to matching start values and linear compensation to compensate for evaporation effects. Control measurements show contributions of unspecific interactions (not shown), complicating data analysis and preventing extraction of the exact association rates of GPX4 binding to the respective nanodiscs or nanodiscs-inhibitor mixtures.

### Lipid Overlay assay

To identify the lipids and sphingolipids that bind to GPX4, Membrane Lipid Strips (Echelon Biosciences S-6002) and Sphingo-Strips (Echelon Biosciences S-6000) were used. The membrane was blocked 1 hour at room temperature using TBS-T + 3% fatty acid-free BSA and then incubated during 2.5 hours at room temperature with 1 µg/mL (50 nM) GPX4 protein (SeLENOZYME). To evaluate ATM, the protein was preincubated with 100 µM ATM for 30 min at room temperature before being added to the strips. For detection, monoclonal anti GPX4 (67763-1-lg, 1:1000, Proteintech) was used as primary antibody, followed by a secondary anti mouse (1:10000, Southern Biotech, 1030-05) and by Clarity™ Western ECL (BioRad) detection.

### Molecular Docking

To obtain information about the binding interactions of aurothiomalate with both the active and allosteric GPX4 sites, we performed molecular docking using AutoDock Vina 1.2.0^65^. The crystal structure of GPX4 (PDB ID: 6HKQ) was obtained from the Protein Data Bank, while the structure of aurothiomalate was obtained from PubChem Compound and optimized using ORCA 5.0 software^66^ at the density functional theory level (DFT) with the wB97x functional^67,68^ and def2-TZVP basis set. The optimization process involved default convergence criteria, followed by geometry optimization and frequency calculations to confirm the stability of the optimized structure.

For the preparation of the target protein, polar hydrogens were added, and protein atoms were assigned Kollman united-atom partial from the AMBER force field (implemented in AutoDock Tools v1.5.7). Ionizable residues were kept at their default protonation states at pH 7.0. For selenocysteine and its selenenic acid form, charges were obtained from the work by Messias *et al*^69^. For the ligand, nonpolar hydrogens were merged and Gasteiger-Marsili charges were computed prior to PDBQT conversion. AutoDock Tools 1.5.7^70^ were used to prepare both the protein and the ligand.

As the 6HKQ crystal structure has an inhibitor (ML162) in the active site^71^, a re-docking procedure was carried out to validate and evaluate the reproducibility of the docking procedure, ensuring it could reproduce the structural conformation of the co-crystallized ligand at the active site of GPX4. The root mean square deviation (RMSD) between the docked and co-crystallized poses of ML162 was calculated to validate the docking accuracy. The docking grid had dimensions of 24 Å × 24 Å × 24 Å with a spacing of 0.375 Å, centered at coordinates x = -23.394, y = 9.887, and z = 1.856. The same grid box and spacing were applied for the allosteric sites. An energy range of 3, exhaustiveness of 8, and up to 10 poses per ligand were adopted for all docking simulations. The protein structure was kept rigid, while the ligand was fully flexible during docking and the refinement. AutoDock Vina reports a docking score from an empirical scoring function that approximates intermolecular interactions (van der Waals, hydrogen bonding, electrostatics, and desolvation) to a rank poses^65^. These values are used here to compare and rank candidate poses within this study. While not absolutely accurate, values reported allowed ranking configurations by energy. The top 10 ligand binding ddates were ranked according to their binding affinities.

### Molecular Dynamics

To study the influence of thiomalate binding on GPX4, molecular dynamics (MD) simulations were performed on two systems: GPX4 without the ligand and GPX4 with thiomalate covalently bound to the U46 residue via a restraint in the Se-S bond. The MD simulations were performed using the X-ray structure of GPX4 (PDB ID: 6HKQ, resolution 1.54 Å)^71^. Protonation states of amino acids were assigned to physiological pH (i.e., Asp and Glu negatively charged, and Lys and Arg positively charged), with all solvent-exposed His residues were protonated at the δ-N atom. The initial structures were solvated in a 12 Å octahedral box using the TIP3P model for water molecules^72^. The system’s total charge was neutralized using Amber’s uniformly charged plasma^73^. Parameters for all residues were taken from the AMBER ff14SB force field^74^ except for the selenocysteine, which used a novel set of Lennard-Jones parameters developed by Estrin and coworkers^69^. These parameters were derived to reproduce solute-water radial pair distribution functions g(r) (RDF) from ab initio molecular dynamics simulations.

For thiomalate, the force field was developed using standard protocols. Parameters were calculated from a DFT calculation using the PBE0 functional^75^ and def2-TZVP basis set. Partial atomic charges were obtained using the Restrained Electrostatic Potential Procedure (RESP)^76^.

Periodic boundary conditions with a 10 Å cutoff and the particle mesh Ewald (PME) summation method for treating electrostatic interactions were applied. Covalent bonds involving hydrogen atoms were restrained at their equilibrium distance using the SHAKE algorithm^77^, which allowed a 2-fs time step for integrating Newton’s equations. Temperature and volume were kept constant with the Langevin thermostat (collision frequency of 2.0 ps⁻¹), and the Berendsen barostat was used to regulate pressure conditions^78^, as implemented in the AMBER20 package^73^.

The equilibration protocol consisted of: (i) 25,000 steps of steepest descent energy optimization followed by 25,000 steps of conjugate gradient energy optimization to prevent close contacts; (ii) slow heating of the whole system (protein and solvation box) from 0 to 300 K for 2 ns at constant volume (iii) six rounds of equilibration at constant pressure (NPT ensemble) at 300 K and 1 atm; (iv) molecular dynamics simulation for approximately 600 ns in the NPT ensemble using the Berendsen barostat and Langevin thermostat, with a 2 fs timestep and final temperature of 300 K. A distance restraint was applied to model the S-Se bond, in which a penalty was applied at distances larger than 3 Å.

All structures were stable during the simulation timescale

### Inductively coupled plasma mass spectrometry (ICP-MS)

Glass vials equipped with teflon septa were cleaned with nitric acid 65% (VWR, Suprapur, 1.00441.0250), washed with ultrapure water (Sigma-Aldrich, 1012620500) and dried. Cells were harvested and washed twice with 1× PBS. Cells were then counted using an automated cell counter (Entek) and transferred in 200 µL 1× PBS or ultrapure water to the cleaned glass vials. The same volume of 1× PBS or ultrapure water was transferred into separate vials for the background subtraction, at least in duplicate per experiment. For tissue samples, a small piece of about 1 mm^3^ was transferred into a clean pre-weighed vial. Samples were lyophilised using a freeze dryer (CHRIST, 2-4 LDplus). Vials with tissue samples were weighed subsequently to determine the tissue dry weight. Samples were then mixed with nitric acid 65% and heated at 80 °C overnight in the same glass vials closed with a lid carrying a teflon septum. Samples were then cooled to room temperature and diluted with ultrapure water to a final concentration of 0.475 N nitric acid and transferred to metal-free centrifuge vials (VWR, 89049-172) for subsequent mass spectrometry analyses. Amounts of metals were measured using an Agilent 7900 ICP-QMS in low-resolution mode, taking natural isotope distribution into account. Sample introduction was achieved with a micro-nebulizer (MicroMist, 0.2 mL/min) through a Scott spray chamber. Isotopes were measured using a collision-reaction interface with helium gas (5 mL/min) to remove polyatomic interferences. Scandium and indium internal standards were injected after inline mixing with the samples to control the absence of signal drift and matrix effects. A mix of certified standards was measured at concentrations spanning those of the samples to convert count measurements to concentrations in the solution. Values were normalised against tissue dry weight.

## REFERENCES

1 Berndt, C. et al. Ferroptosis in health and disease. Redox Biol 75, 103211 (2024). 10.1016/j.redox.2024.103211

2 Hangauer, M. J. et al. Drug-tolerant persister cancer cells are vulnerable to GPX4 inhibition. Nature 551, 247–250 (2017). 10.1038/nature24297

3 Kalkavan, H. et al. Sublethal cytochrome c release generates drug-tolerant persister cells. Cell 185, 3356–3374.e3322 (2022). 10.1016/j.cell.2022.07.025

4 Lei, G. et al. BRCA1-Mediated Dual Regulation of Ferroptosis Exposes a Vulnerability to GPX4 and PARP Co-Inhibition in BRCA1-Deficient Cancers. Cancer Discov 14, 1476–1495 (2024). 10.1158/2159-8290.Cd-23-1220

5 Alborzinia, H. et al. LRP8-mediated selenocysteine uptake is a targetable vulnerability in MYCN-amplified neuroblastoma. EMBO Mol Med 15, e18014 (2023). 10.15252/emmm.202318014

6 Alborzinia, H. et al. MYCN mediates cysteine addiction and sensitizes neuroblastoma to ferroptosis. Nat Cancer 3, 471–485 (2022). 10.1038/s43018-022-00355-4

7 Mai, T. T. et al. Salinomycin kills cancer stem cells by sequestering iron in lysosomes. Nat Chem 9, 1025–1033 (2017). 10.1038/nchem.2778

8 Caneque, T. et al. Activation of lysosomal iron triggers ferroptosis in cancer. Nature 642, 492–500 (2025). 10.1038/s41586-025-08974-4

9 Soula, M. et al. Metabolic determinants of cancer cell sensitivity to canonical ferroptosis inducers. Nat Chem Biol 16, 1351–1360 (2020). 10.1038/s41589-020-0613-y

10 Cheng, Q. & Arner, E. S. Selenocysteine Insertion at a Predefined UAG Codon in a Release Factor 1 (RF1)-depleted Escherichia coli Host Strain Bypasses Species Barriers in Recombinant Selenoprotein Translation. J Biol Chem 292, 5476–5487 (2017). 10.1074/jbc.M117.776310

11 Cheng, Q. A., E. S. J. Overexpression of recombinant selenoproteins in E. coli. Methods in Molecular Biology 1661, 231–240 (2018). 10.1007/978-1-4939-7258-6_17

12 Cheff, D. M., et al. The ferroptosis inducing compounds RSL3 and ML162 are not direct inhibitors of GPX4 but of TXNRD1. Redox Biol 62, 102703 (2023). 10.1016/j.redox.2023.102703

13 DeAngelo, S. L. et al. Recharacterization of the Tumor Suppressive Mechanism of RSL3 Identifies the Selenoproteome as a Druggable Pathway in Colorectal Cancer. Cancer Res 85, 2788–2804 (2025). 10.1158/0008-5472.CAN-24-3478

14 Liu, H. et al. Small-molecule allosteric inhibitors of GPX4. Cell Chem Biol 29, 1680–1693 e1689 (2022). 10.1016/j.chembiol.2022.11.003

15 Cozza, G. et al. Glutathione peroxidase 4-catalyzed reduction of lipid hydroperoxides in membranes: The polar head of membrane phospholipids binds the enzyme and addresses the fatty acid hydroperoxide group toward the redox center. Free Radic Biol Med 112, 1–11 (2017). 10.1016/j.freeradbiomed.2017.07.010

16 Boike, L., Henning, N. J. & Nomura, D. K. Advances in covalent drug discovery. Nat Rev Drug Discov 21, 881–898 (2022). 10.1038/s41573-022-00542-z

17 Eaton, J. K. et al. Selective covalent targeting of GPX4 using masked nitrile-oxide electrophiles. Nat Chem Biol 16, 497–506 (2020). 10.1038/s41589-020-0501-5

18 Wardman, P. & Candeias, L. P. Fenton chemistry: an introduction. Radiat Res 145, 523–531 (1996).

19 Dixon, S. J. et al. Ferroptosis: an iron-dependent form of nonapoptotic cell death. Cell 149, 1060–1072 (2012). 10.1016/j.cell.2012.03.042

20 Xiao, K. et al. Pro-oxidant response and accelerated ferroptosis caused by synergetic Au(I) release in hypercarbon-centered gold(I) cluster prodrugs. Nat Commun 13, 4669 (2022). 10.1038/s41467-022-32474-y

21 Adzavon, K. P., Zhao, W., He, X. & Sheng, W. Ferroptosis resistance in cancer cells: nanoparticles for combination therapy as a solution. Front Pharmacol 15, 1416382 (2024). 10.3389/fphar.2024.1416382

22 Lu, Y. et al. MYCN mediates TFRC-dependent ferroptosis and reveals vulnerabilities in neuroblastoma. Cell Death Dis 12, 511 (2021). 10.1038/s41419-021-03790-w

23 Omata, Y. et al. Sublethal concentrations of diverse gold compounds inhibit mammalian cytosolic thioredoxin reductase (TrxR1). Toxicol In Vitro 20, 882–890 (2006). 10.1016/j.tiv.2006.01.012

24 Stafford, W. C., et al. Irreversible inhibition of cytosolic thioredoxin reductase 1 as a mechanistic basis for anticancer therapy. Sci Transl Med 10 (2018). 10.1126/scitranslmed.aaf7444

25 Samarin, J. et al. Low level of antioxidant capacity biomarkers but not target overexpression predicts vulnerability to ROS-inducing drugs. Redox Biol 62, 102639 (2023). 10.1016/j.redox.2023.102639

26 Doll, S. et al. FSP1 is a glutathione-independent ferroptosis suppressor. Nature 575, 693–698 (2019). 10.1038/s41586-019-1707-0

27 Bersuker, K. et al. The Coti oxidoreductase FSP1 acts parallel to GPX4 to inhibit ferroptosis. Nature 575, 688–692 (2019). 10.1038/s41586-019-1705-2

28 Liu, Y. et al. The 5-Lipoxygenase Inhibitor Zileuton Confers Neuroprotection against Glutamate Oxidative Damage by Inhibiting Ferroptosis. Biol Pharm Bull 38, 1234–1239 (2015). 10.1248/bpb.b15-00048

29 Yang, W. S. et al. Regulation of ferroptotic cancer cell death by GPX4. Cell 156, 317–331 (2014). 10.1016/j.cell.2013.12.010

30 Chen, Z. et al. PRDX6 contributes to selenocysteine metabolism and ferroptosis resistance. Mol Cell 84, 4645–4659.e4649 (2024). 10.1016/j.molcel.2024.10.027

31 Ito, J. et al. PRDX6 dictates ferroptosis sensitivity by directing cellular selenium utilization. Mol Cell 84, 4629–4644 e4629 (2024). 10.1016/j.molcel.2024.10.028

32 Cui, S. et al. Identification of hyperoxidized PRDX3 as a ferroptosis marker reveals ferroptotic damage in chronic liver diseases. Mol Cell 83, 3931–3939 e3935 (2023). 10.1016/j.molcel.2023.09.025

33 Lawson, K. J. D., Christopher J.; Fyfe, David A. The uptake and subcellular distribution of gold in rat liver cells aoer in vivo administration of sodium aurothiomalate. Biochemical Pharmacology 26, 2417–2426 (1977). 10.1016/0006-2952(77)90451-8

34 Kaufmann, K. B. et al. A stemness screen reveals C3orf54/INKA1 as a promoter of human leukemia stem cell latency. Blood 133, 2198–2211 (2019). 10.1182/blood-2018-10-881441

35 Lechman, E. R. et al. miR-126 Regulates Distinct Self-Renewal Outcomes in Normal and Malignant Hematopoietic Stem Cells. Cancer Cell 29, 214–228 (2016). 10.1016/j.ccell.2015.12.011

36 Vučković, A. M. et al. Inactivation of the glutathione peroxidase GPx4 by the ferroptosis-inducing molecule RSL3 requires the adaptor protein 14-3-3ε. FEBS LeE 594, 611–624 (2020). 10.1002/1873-3468.13631

37 Messias, A. et al. Comparing thiol and selenol reactivity towards peroxynitrite by computer simulation. Redox Biochemistry and Chemistry 9 (2024). 10.1016/j.rbc.2024.100035

38 Reich, H. J. & Hondal, R. J. Why Nature Chose Selenium. ACS Chem Biol 11, 821–841 (2016). 10.1021/acschembio.6b00031

39 Stuhlmeier, K. M. The anti-rheumatic gold salt aurothiomalate suppresses interleukin-1beta-induced hyaluronan accumulation by blocking HAS1 transcription and by acting as a COX-2 transcriptional repressor. J Biol Chem 282, 2250–2258 (2007). 10.1074/jbc.M605011200

40 Mertens, R. T., Gukathasan, S., Arojojoye, A. S., Olelewe, C. & Awuah, S. G. Next Generation Gold Drugs and Probes: Chemistry and Biomedical Applications. Chem Rev 123, 6612–6667 (2023). 10.1021/acs.chemrev.2c00649

41 Shen, S., Shen, J., Luo, Z., Wang, F. & Min, J. Molecular mechanisms and clinical implications of the gold drug auranofin. CoordinaGon Chemistry Reviews 493, 215323 (2023). 10.1016/j.ccr.2023.215323

42 Liu, H. et al. Characterization of a patient-derived variant of GPX4 for precision therapy. Nat Chem Biol 18, 91–100 (2022). 10.1038/s41589-021-00915-2

43 Darabi, F. et al. Reactions of model proteins with aurothiomalate, a clinically established gold(I) drug: The comparison with auranofin. J Inorg Biochem 149, 102–107 (2015). 10.1016/j.jinorgbio.2015.03.013

44 Laib, J. E. et al. Formation and characterization of aurothioneins: Au,Zn,Cd-thionein, Au,Cd-thionein, and (thiomalato-Au)chi-thionein. Biochemistry 24, 1977–1986 (1985). 10.1021/bi00329a027

45 Troisi, R., Galardo, F., Messori, L., Sica, F. & Merlino, A. The X-ray structure of the adduct formed upon reaction of aurothiomalate with apo-transferrin: gold binding sites and a unique transferrin structure along the apo/holo transition pathway. Inorganic Chemistry FronGers 12, 2627–2637 (2025). 10.1039/d4qi03184a

46 Zhang, ti. et al. Gold drugs as colistin adjuvants in the fight against MCR-1 producing bacteria. J Biol Inorg Chem 28, 225–234 (2023). 10.1007/s00775-022-01983-y

47 Erdogan, E. et al. Aurothiomalate inhibits transformed growth by targeting the PB1 domain of protein kinase Ciota. J Biol Chem 281, 28450–28459 (2006). 10.1074/jbc.M606054200

48 Stolwijk, J. M., Falls-Hubert, K. C., Searby, C. C., Wagner, B. A. & Buettner, G. R. Simultaneous detection of the enzyme activities of GPx1 and GPx4 guide optimization of selenium in cell biological experiments. Redox Biol 32, 101518 (2020). 10.1016/j.redox.2020.101518

49 Ingold, I. et al. Selenium Utilization by GPX4 Is Required to Prevent Hydroperoxide-Induced Ferroptosis. Cell 172, 409–422 e421 (2018). 10.1016/j.cell.2017.11.048

50 Conrad, M. & Proneth, B. Selenium: Tracing Another Essential Element of Ferroptotic Cell Death. Cell Chem Biol 27, 409–419 (2020). 10.1016/j.chembiol.2020.03.012

51 Lothrop, A. P., Ruggles, E. L. & Hondal, R. J. No selenium required: reactions catalyzed by mammalian thioredoxin reductase that are independent of a selenocysteine residue. Biochemistry 48, 6213–6223 (2009). 10.1021/bi802146w

52 Du, Y., Zhang, H., Lu, J. & Holmgren, A. Glutathione and glutaredoxin act as a backup of human thioredoxin reductase 1 to reduce thioredoxin 1 preventing cell death by aurothioglucose. J Biol Chem 287, 38210–38219 (2012). 10.1074/jbc.M112.392225

53 Gencheva, R. & Arner, E. S. J. Thioredoxin Reductase Inhibition for Cancer Therapy. Annu Rev Pharmacol Toxicol 62, 177–196 (2022). 10.1146/annurev-pharmtox-052220-102509

54 Marti-Andres, P. et al. TRP14 is the rate-limiting enzyme for intracellular cystine reduction and regulates proteome cysteinylation. EMBO J 43, 2789–2812 (2024). 10.1038/s44318-024-00117-1

55 Mansfield, A. S., et al. Phase I dose escalation study of the PKCι inhibitor aurothiomalate for advanced non-small-cell lung cancer, ovarian cancer, and pancreatic cancer. AnGcancer Drugs 24, 1079–1083 (2013). 10.1097/cad.0000000000000009

56 Seo, H., Kohlbrand, A. J., Stokes, R. W., Chung, J. & Cohen, S. M. Masking thiol reactivity with thioamide, thiourea, and thiocarbamate-based MBPs. Chem Commun (Camb) 59, 2283–2286 (2023). 10.1039/d2cc06596g

57 Nakamura, T. et al. A tangible method to assess native ferroptosis suppressor activity. Cell Rep Methods 4, 100710 (2024). 10.1016/j.crmeth.2024.100710

58 Arner, E. S. J. & Schmidt, E. E. Unresolved questions regarding cellular cysteine sources and their possible relationships to ferroptosis. Adv Cancer Res 162, 1–44 (2024). 10.1016/bs.acr.2024.04.001

59 Sies, H., Mailloux, R. J. & Jakob, U. Fundamentals of redox regulation in biology. Nat Rev Mol Cell Biol 25, 701–719 (2024). 10.1038/s41580-024-00730-2

60 Berndt, C., Lillig, C. H. & Flohé, L. Redox regulation by glutathione needs enzymes. Front Pharmacol 5, 168 (2014). 10.3389/fphar.2014.00168

61 Klaus, B. & Reisenauer, S. An end to end workflow for differential gene expression using Affymetrix microarrays. F1000Res 5, 1384 (2016). 10.12688/f1000research.8967.2

62 Demichev, V., Messner, C. B., Vernardis, S. I., Lilley, K. S. & Ralser, M. DIA-NN: neural networks and interference correction enable deep proteome coverage in high throughput. Nat Methods 17, 41–44 (2020). 10.1038/s41592-019-0638-x

63 Pang, Z., et al. Using MetaboAnalyst 5.0 for LC-HRMS spectra processing, multi-omics integration and covariate adjustment of global metabolomics data. Nat Protoc 17, 1735–1761 (2022). 10.1038/s41596-022-00710-w

64 Niesen, F. H., Berglund, H. & Vedadi, M. The use of differential scanning fluorimetry to detect ligand interactions that promote protein stability. Nat Protoc 2, 2212–2221 (2007). 10.1038/nprot.2007.321

65 Eberhardt, J., Santos-Martins, D., Tillack, A. F. & Forli, S. AutoDock Vina 1.2.0: New Docking Methods, Expanded Force Field, and Python Bindings. J Chem Inf Model 61, 3891–3898 (2021). 10.1021/acs.jcim.1c00203

66 Neese, F. Sooware update: The ORCA program system—Version 5.0. WIREs ComputaGonal Molecular Science 12 (2022). 10.1002/wcms.1606

67 Chai, J. D. & Head-Gordon, M. Long-range corrected hybrid density functionals with damped atom-atom dispersion corrections. Phys Chem Chem Phys 10, 6615–6620 (2008). 10.1039/b810189b

68 Grimme, S. Semiempirical GGA-type density functional constructed with a long-range dispersion correction. J Comput Chem 27, 1787–1799 (2006). 10.1002/jcc.20495

69 Pedron, F. N., Messias, A., Zeida, A., Roitberg, A. E. & Estrin, D. A. Novel Lennard-Jones Parameters for Cysteine and Selenocysteine in the AMBER Force Field. J Chem Inf Model 63, 595–604 (2023). 10.1021/acs.jcim.2c01104

70 Morris, G. M., et al. AutoDock4 and AutoDockTools4: Automated docking with selective receptor flexibility. J Comput Chem 30, 2785–2791 (2009). 10.1002/jcc.21256

71 Moosmayer, D., et al. Crystal structures of the selenoprotein glutathione peroxidase 4 in its apo form and in complex with the covalently bound inhibitor ML162. Acta Crystallogr D Struct Biol 77, 237–248 (2021). 10.1107/s2059798320016125

72 Jorgensen, W. L., Chandrasekhar, J., Madura, J. D., Impey, R. W. & Klein, M. L. Comparison of Simple Potential Functions for Simulating Liquid Water. J Chem Phys 79, 926–935 (1983). Doi 10.1063/1.445869

73 Case, D. A., et al. AmberTools. J Chem Inf Model 63, 6183–6191 (2023). 10.1021/acs.jcim.3c01153

74 Lindorff-Larsen, K., et al. Improved side-chain torsion potentials for the Amber ff99SB protein force field. Proteins 78, 1950–1958 (2010). 10.1002/prot.22711

75 Adamo, C. C., M.; Barone, V. An accurate density-functional method for the study of magnetic properties: the PBE0 model. Journal of Molecular Structure: THEOCHEM 493, 145–157 (1999). 10.1016/S0166-1280(99)00235-3

76 Woods, R. J. C., R. Restrained electrostatic potential atomic partial charges for condensed-phase simulations of carbohydrates. Journal of Molecular Structure: THEOCHEM 527, 149–156 (2000). 10.1016/S0166-1280(00)00487-5

77 Ryckaert, J.-P., Ciccow, G. & Berendsen, H. Numerical integration of the Cartesian Equations of Motion of a System with Constraints: Molecular Dynamics of n-Alkanes. JOURNAL OF COMPUTATIONAL PHYSICS 23, 327–341 (1977). 10.1016/0021-9991(77)90098-5

78 Berendsen, H. J. C., Postma, J. P. M., Vangunsteren, W. F., Dinola, A. & Haak, J. R. Molecular-Dynamics with Coupling to an External Bath. J Chem Phys 81, 3684–3690 (1984). Doi 10.1063/1.448118

