## Supplementary Material for "Sodium Aurothiomalate Induces Ferroptosis by Targeting GPX4 via Gold-Dependent Thiomalate Covalent Modification"

### Supplementary Figure 1:

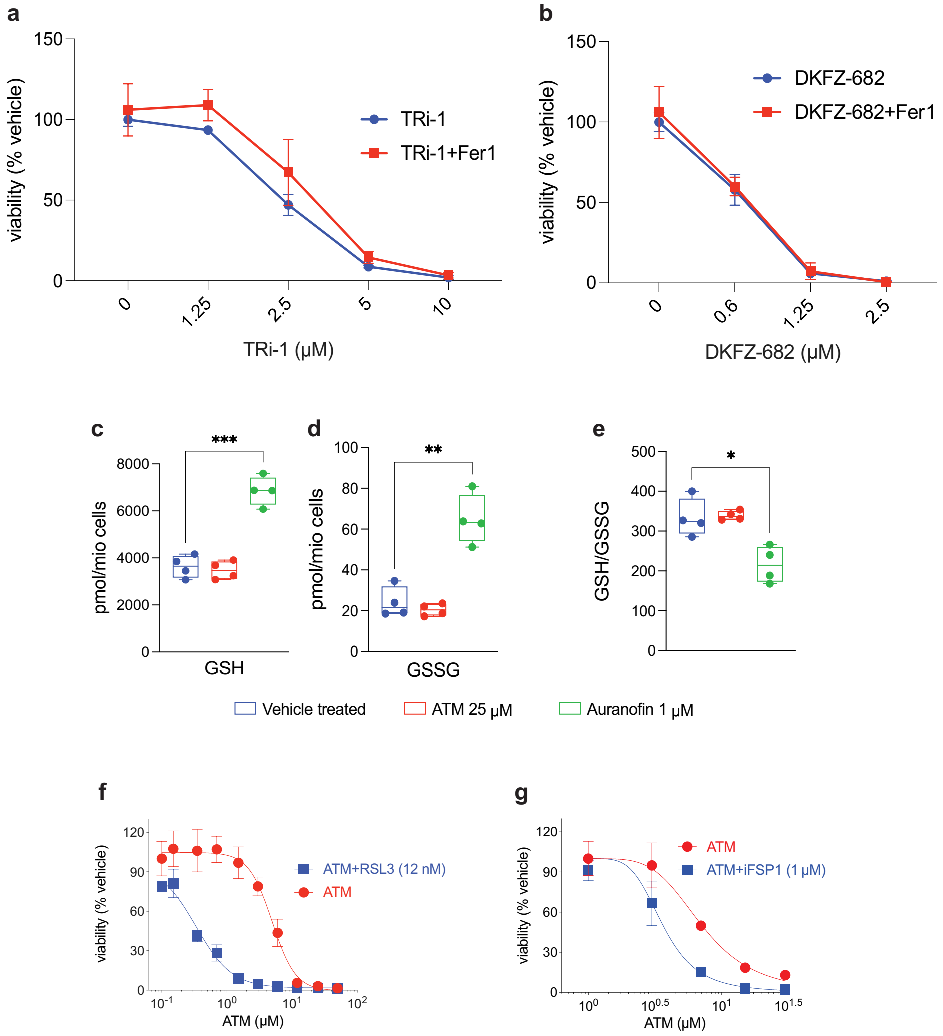


**Supp Fig. 1: a, b)** SK-N-DZ cells treated with TXNRD1 inhibitors TRi-1 and DKFZ-682 for 3 days in the presence or absence of Fer1. **c, d, e)** Quantitative analysis of total intracellular glutathione (GSH) levels, glutathione disulfide (GSSG) levels (n = 4), and the GSH/GSSG ratio (n = 4) in SK-N-DZ cells treated for 24 hours with indicated concentrations of ATM or Auranofin. Data are presented as mean ± standard deviation (SD)**. f)** SK-N-DZ cells treated with ATM in combination with a sublethal dose of the GPX4 inhibitor RSL3 for 3 days. **g)** SK-N-DZ cells treated with ATM in combination with a sublethal dose of the FSP1 inhibitor iFSP1 for 3 days.

**Supplementary Figure 2:**

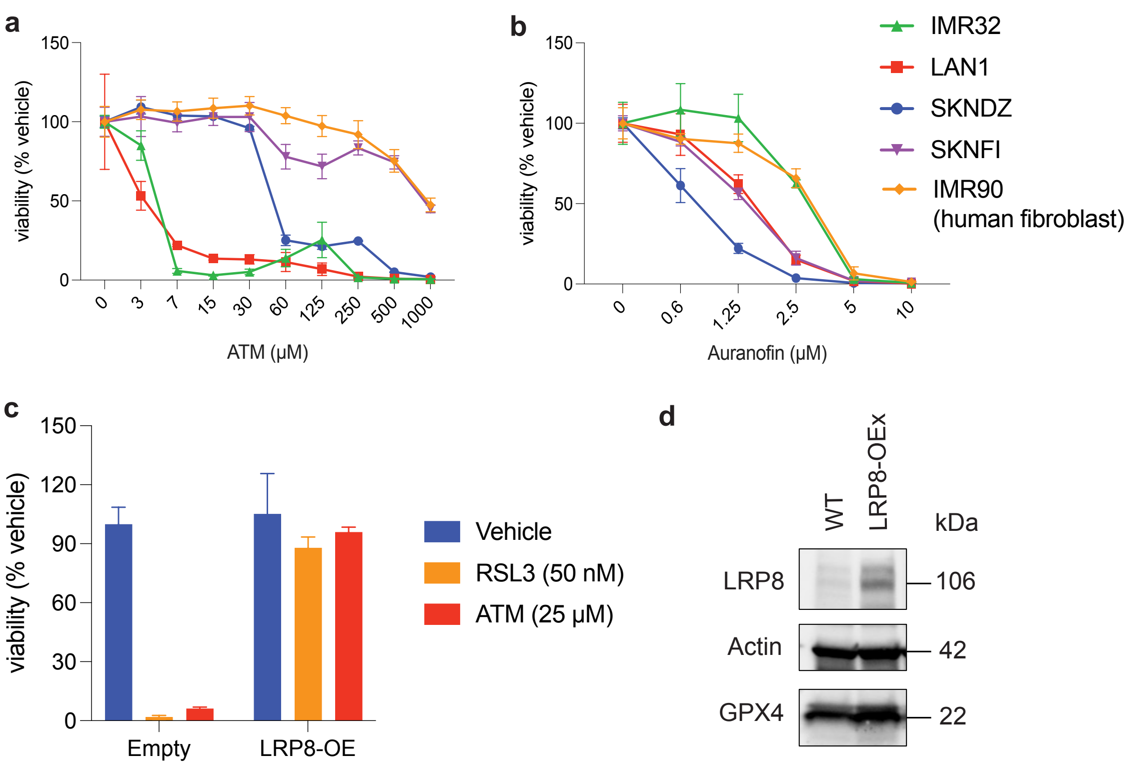


**Suppl. Fig. 2: a-b)** Cellular responses of neuroblastoma cell lines to 72 h of ATM (in **a**) or Auranofin (in **b**) treatment: the experiment includes neuroblastoma cells with *MYCN* amplification (IMR-31,LAN-1, SK-N-DZ), moderate *MYCN* expression (SKNFI), as well as normal human fibroblasts (IMR90). Data represent the mean ± s.e.m.; $n = 3$ samples. **c)** Impact of LRP8 overexpression on SK-N-DZ cell viability analyzed using CellTiter-Glo after treatment with RSL3 or ATM for 3 days (left panel). LRP8 overexpression confirmed by immunoblot analysis (right panel). Data are presented as mean ± SEM. **d)** Western Blot of both WT (left) and LRP8 overexpressing (right) SK-N-DZ cells.

### Supplementary Figure 3:

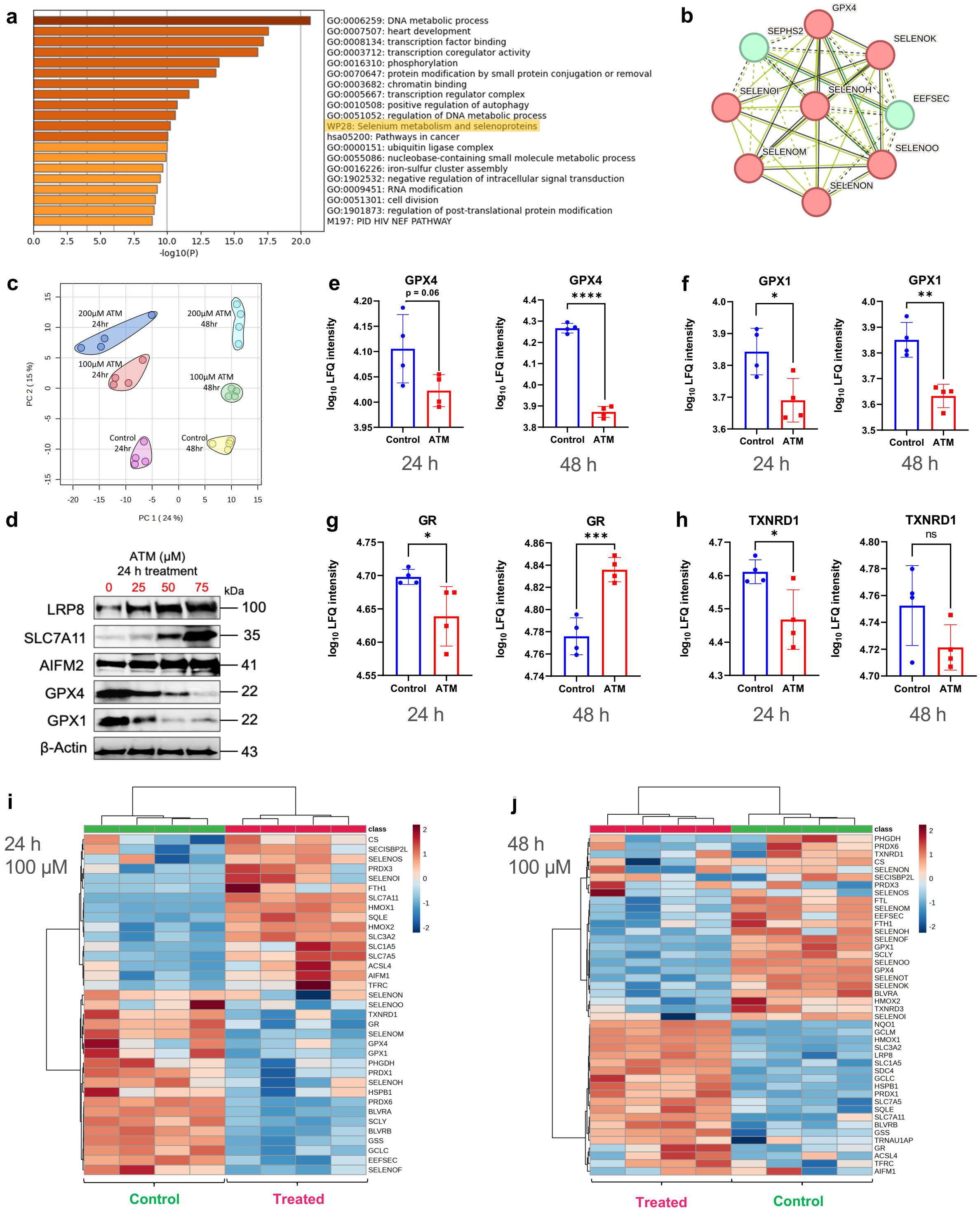

**Suppl. Fig. 3: a)** GO and Pathway analysis of differentially expressed proteins upon ATM treatment (48 hours 200 μM) highlights "selenium metabolism and selenoproteins" (in yellow) as key pathway **b)** Protein-Protein Interaction network of proteins involved in the "Selenium Metabolism and selenoproteins" term. **c)** Proteomics PCA analysis of SK-N-DZ samples treated with two concentrations of ATM (100 and 200 μM), during two different incubation timepoints (24 and 48 hours). e-h) $log_{10}$ LFQ intensity plots comparing protein levels after 24 h (left) or 48 h (right) treatment either with vehicle or 200 μM ATM. **d)** Western blot analysis was performed on SK-N-DZ cells that were either vehicle-treated or treated with the indicated concentrations of ATM for 24 hours. β-actin was used as the loading control. **e-h)** Comparison of protein levels after 24 and 48 hours of 200 μM ATM treatment. ****P < 0.0001 ***P < 0.001; **P < 0.01. *P < 0.05. **e)** GPX4, **f)** GPX1, **g)** GR and **h)** TXNRD1. **i,j)** Protein level changes of ferroptosis markers of SK-N-DZ cells after 24 **(i)** or 48 **(j)** hours of 100 μM ATM in comparison to cells without treatment.

**Supplementary Figure 4:**

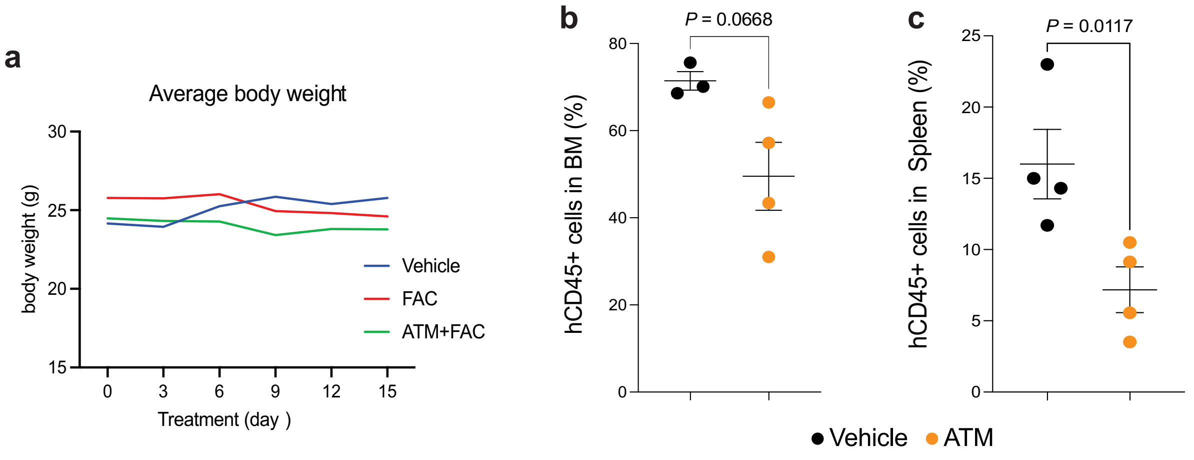


**Suppl. Fig. 4: a)** Body weight of mice treated with the ATM/FAC combination during the treatment period. **b, c)** Percentage of primary AML cells (AML03) in the bone marrow and spleen following treatment with the ATM/FAC combination or vehicle control. Each dot represents the bone marrow or spleen tumor burden in an individual mouse (n = 4). Data are presented as mean ± SEM.

### Supplementary Figure 5:

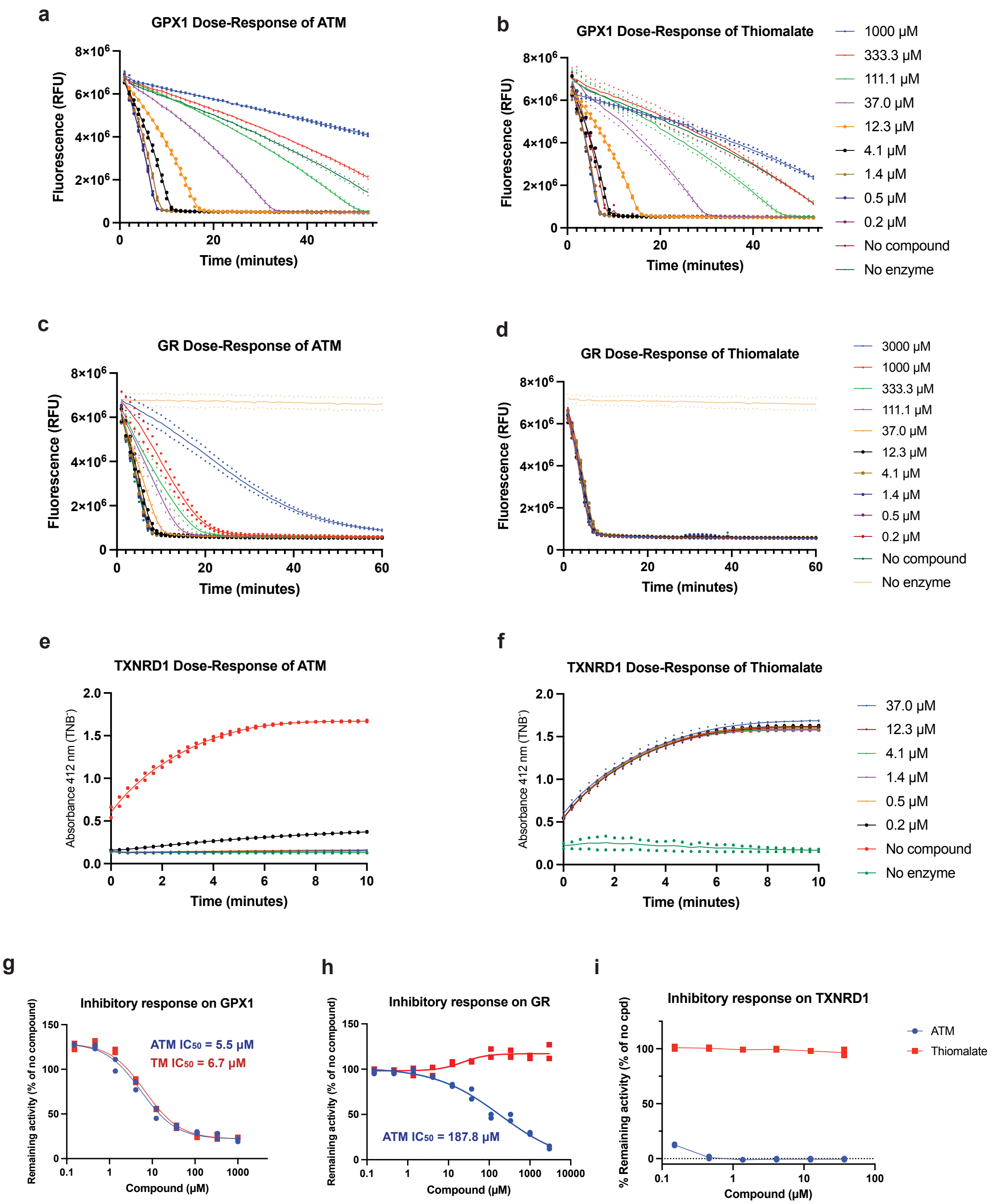


**Suppl. Fig. 5: a-b)** The activity of purified recombinant SeCys-containing human GPX1 was measured using an enzyme coupled assay with glutathione reductase (GR), reduced glutathione (GSH) and NADPH, in the absence and presence of a wide range of ATM (a) or Thiomalate (b) concentrations (0.2-3000 µM). The reaction was initiated by the addition of 0.5 mM (final concentration in well) Cumene hydroperoxide to the reaction mixture and the NADPH fluorescence (λex = 340 nm; λem = 450 nm) was monitored over time, in a similar way as previously described by Cheff et al[1]. **c-d)** The activity of GR was measured in a reaction initiated by the addition of oxidized glutathione (GSSG) and NADPH in Assay Buffer for final concentration 0.4 mM NADPH and 1 mM GSSG, and NADPH fluorescence (λ_ex = 340 nm; λ_em = 450 nm) was monitored over time. The effect of ATM **(c)** and Thiomalate **(d)** was evaluated on GR activity. **e-f)** The activity of TXNRD1 was measured by monitoring $TNB^-$ formation (λ_abs = 412 nm**)** over time in an Assay containing NADPH and using different concentrations of ATM **(e)** or thiomalate **(f)**. The reaction was initiated by the addition of DTNB for final concentration 2.5 mM. **g-h)** Four-parameter dose-response curve fit to n = 2 technical replicates and $IC_{50}$ calculations of ATM and Thiomalate for GPX1 **(g)** and GR **(h)**. **i)** Inhibitory response of ATM and thiomalate on TXNRD1.

### Supplementary Figure 6

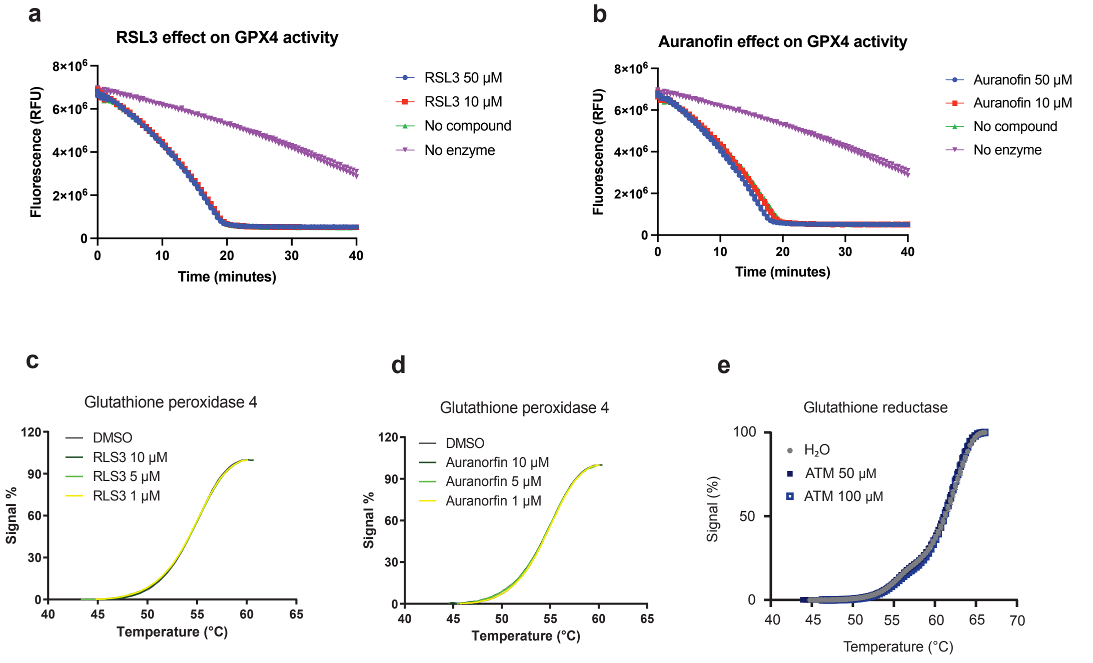


**Suppl. Fig. 6: a-b)** Evaluation of the effect of RSL3 **(a)** and auranofin **(b)** on GPX4 activity. **c-e)** Thermal shift assay was conducted to monitor potential changes in protein thermostability of GPX4 caused by RSL3 **(c)** and auranofin **(d)** treatment. In both cases, there are no changes of the melting temperature upon incubation with the compounds**. e)** Thermal shift assay of Glutathione reductase (GR) shows no changes upon ATM incubation.

**Supplementary Figure 7:**

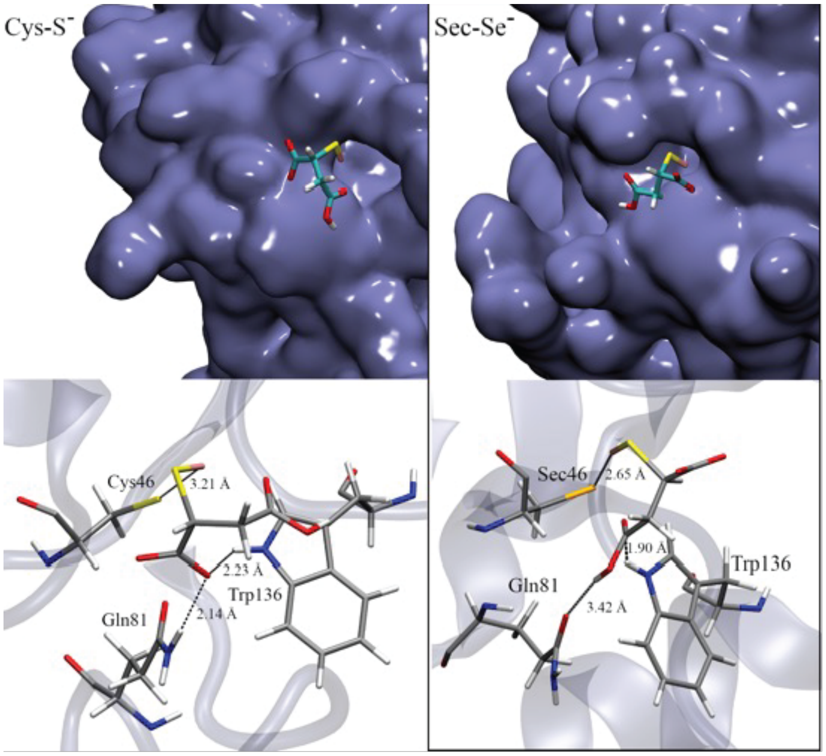


**Suppl. Fig. 7:** Docking results for ATM in the region involving the Sec residue. As observed, other residues (Trp186 and Gln81) may also provide key interactions to ensure the correct inhibitor binding in the groove. A slightly better arrangement is provided for Sec in comparison to that of Cys.

**Supplementary Figure 8:**

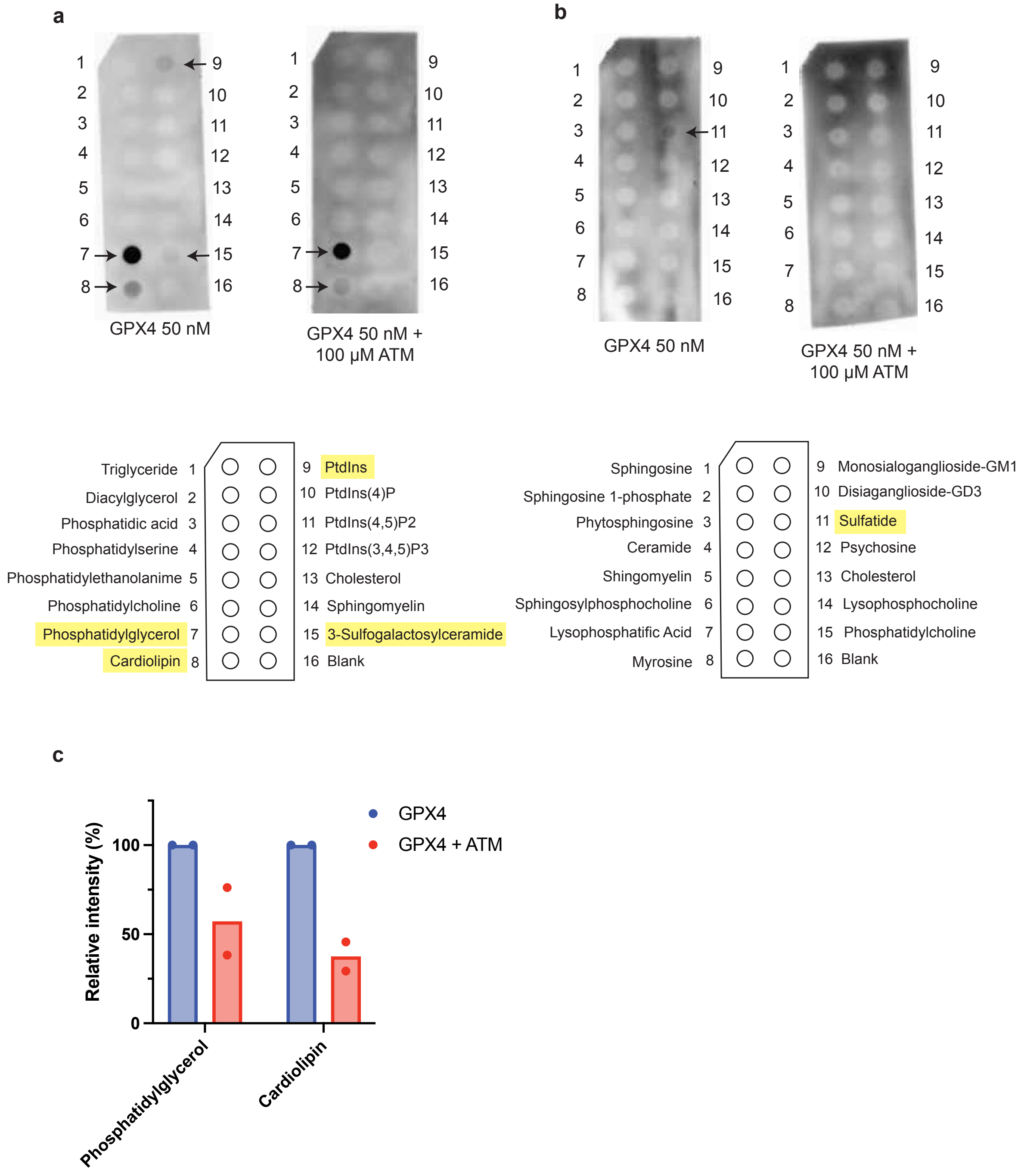


**Suppl. Fig. 8:** Interaction between GPX4 and different lipids and sphingolipids determined using lipid strips. Effect of ATM was determined preincubating the protein 30 min at room temperature before adding it to the membranes. The experiment was performed twice with similar results, the figures correspond to one representative experiment. **a)** When using the lipid strip from Echelon (S-6002), binding of GPX4 was observed mainly to Phosphatidylglycerol and Cardiolipin, as well as low binding interactions with 3-Sulfogalactosylceramide and Phosphatidylinositol. In the presence of ATM those interactions are reduced. **b)** Sphingostrip from Echelon (S-6000) showed low affinity binding to Sulfatide that was affected upon presence of ATM. **c)** Quantification of phosphatidylglycerol and cardiolipin signal intensities from two independent experiments (n = 2). GPX4 intensities in the presence of ATM were normalized to the corresponding values obtained in the absence of ATM for each experiment.

**Supplementary information:**

Methodology computational calculations

In order to investigate the reaction feasibility of Au-S-TML with selenocysteins and cysteins, we conducted a series of electronic structure calculations. The methylated systems methanelthiol (MeSH), methaneselenol (MeSeH), methanesulfenic acid (MeSOH) and methaneselenenic acid (MeSeOH) were selected as simplified models of Cys-SH, Sec-SeH, Cys-SOH and Sec-SeOH respectively. These models were chosen due to their structural simplicity and relevance in mimicking the chemical properties of the amino acid derivatives. To provide a detailed analysis of the higher acidity of MeSeH compared to MeSH and its possible influence on their observed reactivity under physiological conditions, we analyzed the intrinsic reactivity of these compounds towards aurothiomolate Au-S-TML through reactions R1 to R4:

MeSH (aq) + Au-S-TML(aq) → MeS-S-TML (aq) + AuH (aq) (R1)

MeSeH (aq) + Au-S-TML (aq) → MeSe-S-TML (aq) + AuH (aq) (R2)

MeSOH (aq) + Au-S-TML(aq) → MeS-S-TML (aq) + AuOH (aq) (R3)

MeSeOH (aq) + Au-S-TML (aq) → MeSe-S-TML (aq) + AuOH (aq) (R4)

To obtain valid starting structures for further analysis, we studied the reaction of the model systems MeSH, MeSeH, MeSOH and MeSeOH with Au-S-TML (reactions R1 to R4) employing electronic structure calculations in implicit water at the PCM level, using ORCA 5.0. The structures of reactants complex and products complex (i.e. the reactants or products in proximity of each other) of reactions R1, R2, R3 and R4 were optimized in aqueous solution via the PCM approach. The optimization process utilized the range-separated hybrid functional wB97x, along with the def2-TZVP basis set to ensure accurate treatment of the electronic structure.

Frequency calculations were performed in all cases using the harmonic approximation as implemented in ORCA, to confirm the nature of the stationary points and to compute thermochemical properties. All calculations were carried out with default convergence criteria of $10^{-8}$ for energy, $10^{-5}$ for gradient, and $10^{-5}$ for displacement. Additionally, tight integration grids (Grid4 in ORCA, corresponding to a Lebedev grid with 434 angular points) were used to improve the accuracy of the results. The resulting structures and energies were analyzed to determine the Gibbs free energy changes ($\Delta G_{reac}$) for the reactions. All input and output files generated during the calculations are provided in the Supplementary Material

The Gibbs free energy ($\Delta G_{react}$) for the reactions of methanethiol (MeSH), methaneselenol (MeSeH), methanesulfenic acid (MeSOH), and methaneselenenic acid (MeSeOH) with aurothiomolate (Au-S-TML) are summarized in the following table:

**Supplementary table 1:**

| Reaction | $\Delta G_{react}$ (kcal/mol) |
|---|---|
| R1 | -3.18 |
| R2 | -4.40 |
| R3 | -5.50 |
| R4 | -5.72 |

The results indicate that the reactions of MeSeH and MeSeOH with Au-S-TML are more favorable compared to their sulfur analogs. This trend can be attributed to the higher acidity and lower bond dissociation energy of selenium-containing compounds compared to sulfur-containing compounds[2,3].

In the context of these reactions, aurothiomalate acts as an acceptor for sulfur and selenium nucleophiles.

The higher acidity of MeSeH is reflected in the more negative $\Delta G_{reac}$ for reaction R2 compared to R1. This suggests that MeSeH is more likely to donate a proton and react with Au-S-TML, forming MeSe-S-TML and AuH. The greater stability of the selenolate ion ($MeSe^{-}$) compared to the thiolate ion ($MeS^{-}$) contributes to the observed higher reactivity.

The reactions involving MeSOH and MeSeOH (R3 and R4) show more negative $\Delta G_{reac}$ values than their thiol and selenol counterparts (R1 and R2). The formation of MeS-S-TML and MeSe-S-TML from MeSOH and MeSeOH, respectively, is energetically favorable, with the MeSeOH reaction being the most exergonic. This indicates that selenenic acids are more reactive towards Au-S-TML than sulfenic acids, likely due to the lower bond dissociation energy of the Se-O bond compared to the S-O bond[4].

1 Cheff, D. M. *et al.* The ferroptosis inducing compounds RSL3 and ML162 are not direct inhibitors of GPX4 but of TXNRD1. *Redox Biol* **62**, 102703 (2023). https://doi.org/10.1016/j.redox.2023.102703

2 Messias, A. *et al.* Comparing thiol and selenol reactivity towards peroxynitrite by computer simulation. *Redox Biochemistry and Chemistry* **9** (2024). https://doi.org/10.1016/j.rbc.2024.100035

3 Cardey, B. & Enescu, M. Selenocysteine versus cysteine reactivity: a theoretical study of their oxidation by hydrogen peroxide. *J Phys Chem A* **111**, 673-678 (2007). https://doi.org/10.1021/jp0658445

4 Cardey, B. & Enescu, M. A computational study of thiolate and selenolate oxidation by hydrogen peroxide. *Chemphyschem* **6**, 1175-1180 (2005). https://doi.org/10.1002/cphc.200400568
